# Antidepressant Effects of Lauric Acid in a Corticosterone-Induced Murine Model of Depression: Behavioral and Neurochemical Insights

**DOI:** 10.64898/2026.05.15.725442

**Authors:** Manoel C. de Paulo, Lissiana M.V. Aguiar, Nívea M. Cunha Ferreira, Jordana M. Nascimento, Leonardo Resstel, Carla T. V. Melo

## Abstract

**Background:** Lauric acid (LA) is a medium-chain saturated fatty acid found in several foods, including vegetable oils and seeds. Previous studies have demonstrated that LA exhibits neuroprotective, antioxidant, and anti-inflammatory properties in experimental models of neuropsychiatric disorders. Therefore, the present study aimed to investigate the behavioral and neurochemical effects of LA in a corticosterone-induced murine model of depression.

**Methods:** Male Swiss mice received corticosterone (CORT; 20 mg/kg, subcutaneously) for 23 consecutive days, while the control group received vehicle only. During the last nine days of the experimental protocol, the animals received the respective treatments by oral gavage: LA (10 or 20 mg/kg), fluvoxamine (FLUV; 50 mg/kg), or vehicle, administered 1 hour after CORT injection. One hour after treatment administration, the animals were subjected to the behavioral tests: Forced Swimming Test (FST), Tail Suspension Test (TST), and Open Field Test (OFT). At the end of the experimental protocol, the animals were euthanized, and the prefrontal cortex (PFC), hippocampus (HPC), and striatum (STR) were collected for neurochemical analyses.

**Results:** Chronic CORT treatment significantly increased immobility time in the FST and TST, characterizing depressive-like behavior. Treatment with LA reversed these behavioral alterations, showing an effect similar to that observed in the FLUV-treated group. In the OFT, LA did not promote significant changes in locomotor activity, suggesting the absence of psychostimulant effects. Regarding neurochemical analyses, LA treatment did not reduce malondialdehyde (MDA) or nitrite/nitrate (NO_2_^−^/NO_3_^−^) levels, nor did it alter reduced glutathione (GSH) levels in the evaluated brain regions.

**Conclusion:** The results demonstrated that LA treatment was able to reverse corticosterone-induced behavioral alterations in mice, indicating a potential antidepressant-like effect. Furthermore, the observed effects were not associated with nonspecific locomotor alterations. Although LA did not promote significant changes in the evaluated neurochemical markers, these findings reinforce its potential as a therapeutic agent for depressive disorders and highlight the need for further studies to elucidate its mechanisms of action and possible clinical applicability.

## Introduction

Depression is a common mental disorder that affects the lives of individuals suffering from this disease. It is caused by several factors, including genetic, environmental, and psychological influences. Data from the World Health Organization confirm that women are more affected than men, occurring frequently in individuals of all ages (World Health Organization, 2023).

Symptoms of depression include feelings of sadness, guilt, psychomotor retardation, insomnia and drowsiness, decreased appetite, reduced sexual interest, pain, and diffuse physical symptoms (Ministry of Health, 2023). In addition, there is a substantial relationship with anxiety, which may combine recurrent symptoms, fear of the future, self-criticism, among other manifestations (Miller and Hen, 2015).

The prevalence of depression occurs idiopathically; however, there are numerous risk factors that contribute to the development of this disorder, such as cancer, stroke, medication side effects, and repeated exposure to stressful factors (Panjwani and Li, 2021; Sebestova et al., 2021). Stress is one of the major causative factors of depression (Junior et al., 2021). Furthermore, it has been suggested that hypercortisolism leads to increased neurotoxicity in the hippocampus, resulting in the emergence of depressive symptoms (Holsboer et al., 2000; Sterner and Kalynchuk, 2010; Schoenfeld and Cameron, 2015).

During the depressive process, certain physiological alterations result in dysregulation of the Hypothalamic–Pituitary–Adrenal (HPA) axis, which is responsible for controlling glucocorticoid levels in the bloodstream (Liu et al., 2017; Holsboer, 2000; Gogos et al., 2009). These alterations appear to depend on hippocampal integrity, indicating that individuals with depressive symptoms present reduced hippocampal volume (Lorenzetti et al., 2009), and that reversal of these symptoms occurs through increased neurogenesis promoted by several antidepressants (Liu et al., 2017; Miller and Hen, 2015).

The search for new pharmacological therapeutic options for the treatment of depression is a way to help reduce the growing number of treatment discontinuations. In this context, substances such as lauric acid, which acts through different pathways, emerge as alternatives for the treatment of depression.

Lauric acid (LA), also known as dodecanoic acid, is a medium-chain fatty acid and the main component of coconut oil, also found in human milk, cow’s milk, and goat’s milk (Correia et al., 2020). Studies conducted by Sandapuma et al. (2022), Bansal et al. (2019), and Nakagima & Kunugi (2020) demonstrated the neuroprotective properties of lauric acid in Alzheimer’s disease models by reducing intracellular ARF1 expression, thereby inhibiting APP and β-amyloid secretion. In addition, Barreto (2020) observed the effects of low doses in an experimental mania-like model, promoting increased endogenous GSH, reduced lipid peroxidation and nitrite levels, as well as increased BDNF levels, suggesting its potential as a preventive therapy for bipolar disorder. Furthermore, Ribeiro (2024) demonstrated the effects of lauric acid on depression using a predictive model, in which doses of 10, 20, and 40 mg/kg modulated depressive-like behaviors and improved endogenous GSH levels.

Based on this evidence, the aim of the present study was to evaluate the potential antidepressant effect of lauric acid in mice subjected to a depression model induced by chronic corticosterone administration.

## Materials and Methods

### Animals

Male Swiss mice (25–30 g; age: 4–5 weeks) were used in this study. The animals were maintained at room temperature (24–26°C), with free access to food and water *ad libitum*, and were randomly distributed into the specified experimental groups. Animals were housed under a 12 h light/dark cycle (lights off at 18:00 h). All experiments were conducted between 08:00 and 16:00 h. The experimental procedures were approved by the Ethics Committee of the Department of Physiology and Pharmacology of the Federal University of Ceará (Protocol No. 23/01).

### Drugs

Corticosterone (Sigma®, St. Louis, MO, USA) was dissolved in 0.1% polysorbate (Tween®) 80 (VETEC™, USA) and 0.1% dimethyl sulfoxide (DMSO) (VETEC™, USA), diluted in distilled water, and administered subcutaneously (s.c.) at a dose of 20 mg/kg.

Lauric acid (Sigma®, St. Louis, MO, USA) was dissolved in 0.9% saline solution containing 0.1% Tween® 80 and 0.1% DMSO (VETEC™, USA), and administered intragastrically (oral gavage) at doses of 10 and 20 mg/kg. Fluvoxamine (Abbott®, New Jersey, USA), at a dose of 50 mg/kg, was dissolved in distilled water and administered by oral gavage.

### Experimental Procedure

This study was based on a corticosterone-induced murine model of depression, as previously described by Zhao et al. (2008), in which subcutaneous administration of corticosterone (20 mg/kg/day) for up to five weeks induces depressive-like behaviors in mice.

A total of 90 animals were used and randomly distributed into three independent experimental sets, each consisting of 30 animals divided into five experimental groups (*n* = 6 animals/group), resulting in a final *N* of 18 animals per experimental group:

1. Control (vehicle)
2. Corticosterone (CORT)
3. Corticosterone + lauric acid 10 mg/kg (CORT + LA 10)
4. Corticosterone + lauric acid 20 mg/kg (CORT + LA 20)
5. Corticosterone + fluvoxamine 50 mg/kg (CORT + FLUV 50)

The CORT, CORT + LA 10, CORT + LA 20, and CORT + FLUV 50 groups received corticosterone (20 mg/kg, subcutaneously) for 23 consecutive days, while the control group received vehicle only (0.9% saline solution containing 0.1% Tween® 80).

During the last nine days of the experimental protocol, the animals received the respective treatments by oral gavage: lauric acid (10 or 20 mg/kg), fluvoxamine (50 mg/kg), or vehicle, administered 1 hour after corticosterone injection. One hour after treatment administration, the animals were subjected to the behavioral tests Forced Swimming Test (FST), Tail Suspension Test (TST), and Open Field Test (OFT). At the end of the experiments, the animals were euthanized, and the prefrontal cortex, hippocampus, and striatum were dissected for biochemical analyses related to oxidative stress (Figure 1).

**Fig 1.**
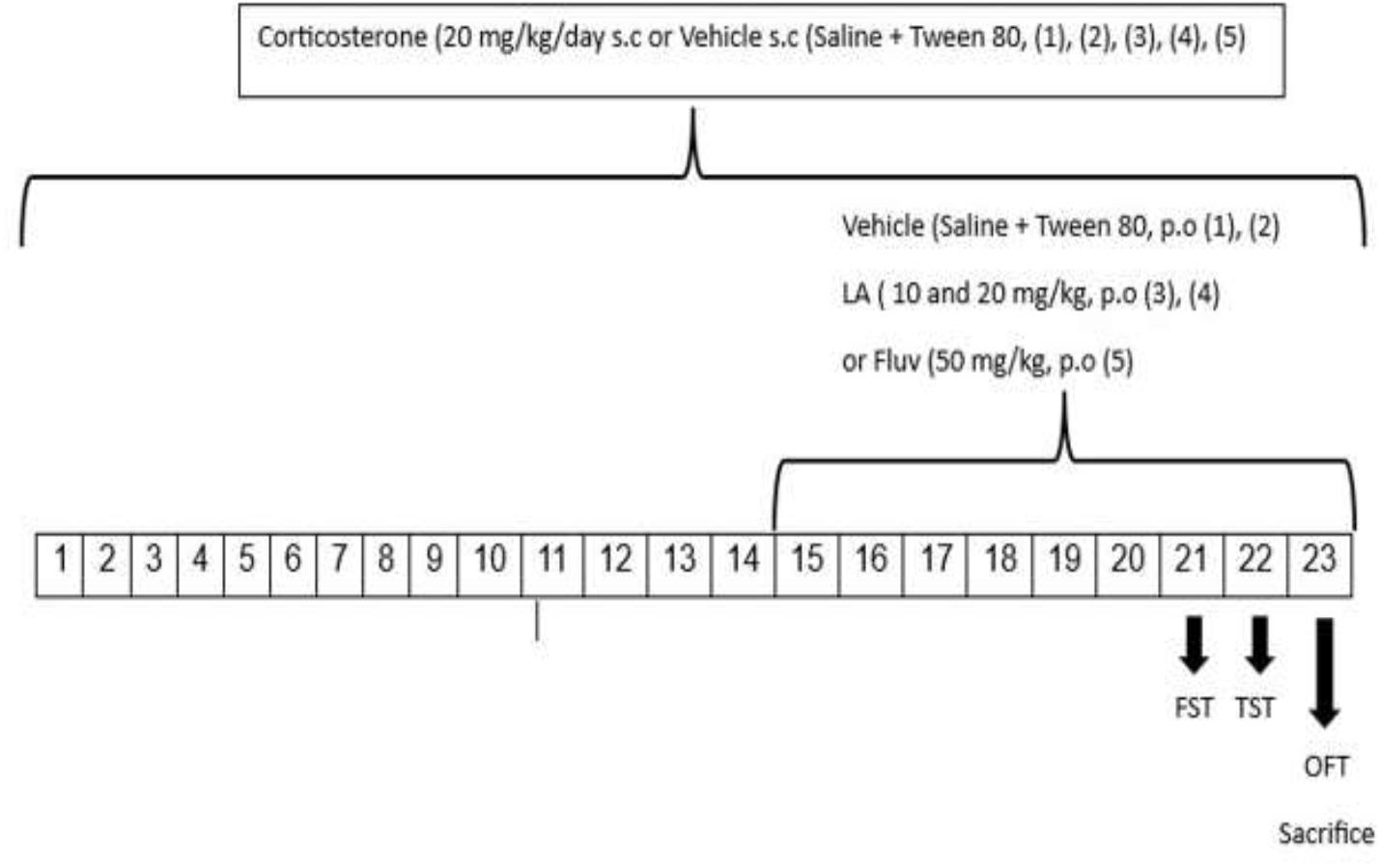
Schematic overview of the experimental design: LA: lauric acid; Fluv: fluvoxamine; Cort: corticosterone; S.C: subcutaneous; P.O: oral gavagem; <u>FST</u> : forced swimming test; <u>TST</u> : tail suspension test; OFT: open field test.

The experimental groups were randomly allocated, and the researcher responsible for handling the animals was blinded to the treatments and identification of the cages containing the animals. The coding procedure was performed by a researcher who had no contact with the animals and did not participate in the analysis of the results. The codes were revealed only after completion of the statistical analyses.

### Behavioral Tests

#### Forced Swimming Test

For this experiment, cylindrical glass tanks containing water at room temperature (25 ± 1 °C) with a depth of 20 cm were used. Sixty minutes after treatment administration, the animals were individually placed in the cylindrical tanks. Immobility time was recorded in seconds over a 5-minute period. Immobility was defined as the time during which the animal remained floating in the water without any movement, except those necessary to keep its nose above the water surface, as described by Porsolt (PORSOLT et al., 1977).

#### Tail Suspension Test

In this experiment, previously described by Steru et al. (1985), 60 minutes after treatment administration, mice were suspended individually by the tail, approximately 1 cm from the tip, using adhesive tape, at a height of 50 cm above the floor. The parameters evaluated were immobility time and latency, measured in seconds during a 5-minute period. All animals were subjected to this behavioral test.

#### Open Field Test

This test, described by Archer (1973), was conducted to evaluate the exploratory activity of the animals. The apparatus consisted of a wooden box with a black floor (30 × 30 × 15 cm). The arena floor was divided into nine equal squares. Sixty minutes after treatment administration, the room was maintained under dim lighting conditions, and a red light was positioned above the apparatus. Each animal was individually and carefully placed in the center of the apparatus, and the following parameters were evaluated over a 5-minute period: the number of squares crossed with all four paws (spontaneous locomotor activity), rearing behavior (vertical exploratory activity), grooming (self-cleaning movements), and freezing time.

### Neurochemical Tests

#### Determination of Lipid Peroxidation by Measurement of Thiobarbituric Acid Reactive Substances (TBARS)

The degree of lipid peroxidation was evaluated according to the method proposed by Draper and Hadley (1990). In this experimental protocol, lipid peroxidation was measured by determining the concentrations of thiobarbituric acid reactive substances (TBARS). Homogenates were prepared from the brain regions of interest: hippocampus, prefrontal cortex, and striatum. The homogenates were prepared in 10% solution of 150 mM phosphate buffer, pH 7.0. Subsequently, 666 µL of 35% perchloric acid was added to the homogenate, followed by centrifugation at 5000 rpm for 10 minutes at 4 °C. The supernatant was then collected, and 250 µL of 1.2% thiobarbituric acid was added. The mixture was incubated in a water bath for 30 minutes at 95 °C. After incubation, the contents were transferred to cuvettes and analyzed in a spectrophotometer at 535 nm. Results were expressed as micrograms (µg) of malondialdehyde (MDA) per gram (g) of tissue.

#### Determination of Nitrite/Nitrate Concentration

To determine nitrite/nitrate production, a standard NaNO_2_ calibration curve was initially prepared. Briefly, 6.9 mg of NaNO_2_ were weighed and dissolved in 10 mL of bidistilled water (stock solution – 10 mM), followed by serial dilutions (10- and 20-fold), resulting in concentrations of 1 mM, 100 µM, 10 µM, 5 µM, 2.5 µM, 1.25 µM, 0.625 µM, and 0.312 µM. A linear equation derived from the standard curve was used to calculate the concentrations of the test samples.

Homogenates from the brain regions were prepared at 10% (1:9) in 150 mM phosphate buffer solution, pH 7.0. The homogenates were then centrifuged at 11,000 rpm for 15 minutes at 4 °C, and the supernatants were collected. Nitric oxide production was measured using the Griess reaction. An aliquot of 500 µL of the supernatant was incubated with 500 µL of Griess reagent (5% phosphoric acid + 1% sulfanilamide diluted in 5% phosphoric acid + 0.1% N-(1-naphthyl)ethylenediamine dihydrochloride [NEED] + distilled water, in a 1:1:1:1 ratio) at room temperature for 10 minutes. Absorbance was measured in a spectrophotometer at 570 nm. Nitrite concentration was determined from the NaNO_2_ standard curve, and results were expressed as micromolar (µM) nitrite/nitrate per gram of tissue, according to Green et al. (1981).

#### Determination of Reduced Glutathione (GSH) Concentration

The determination of GSH concentration is based on the reaction of Ellman’s reagent (5,5’-dithiobis-2-nitrobenzoic acid) with free thiol groups, forming a mixed disulfide and 2-nitro-5-thiobenzoic acid. To determine GSH concentration, homogenates from the brain regions of interest were prepared in 0.02 M EDTA solution. Subsequently, 400 µL of the homogenate were collected and mixed with 500 µL of distilled water and 100 µL of 50% trichloroacetic acid. The mixture was centrifuged at 500 rpm for 15 minutes at 4 °C. Then, 600 µL of the supernatant were collected and added to 846 µL of 0.4 M Tris-HCl solution, pH 8.9, together with 0.01 M DTNB. After 1 minute of reaction, absorbance was measured in a spectrophotometer at 412 nm, according to Sedlak and Lindsay (1988). Reduced glutathione concentration was expressed as µg of GSH per gram of tissue.

## Results

### Antidepressant-Like Effect in the Forced Swimming Test (FST) and Tail Suspension Test (TST)

In the Forced Swimming Test, animals treated with corticosterone showed a significant increase in immobility time when compared to the healthy control group. Animals treated with lauric acid at doses of 10 mg/kg and 20 mg/kg exhibited a significant reduction in immobility time compared to the corticosterone- and vehicle-treated group. A similar result was observed in the group treated with fluvoxamine (Figure 2A).

**Fig 2:**
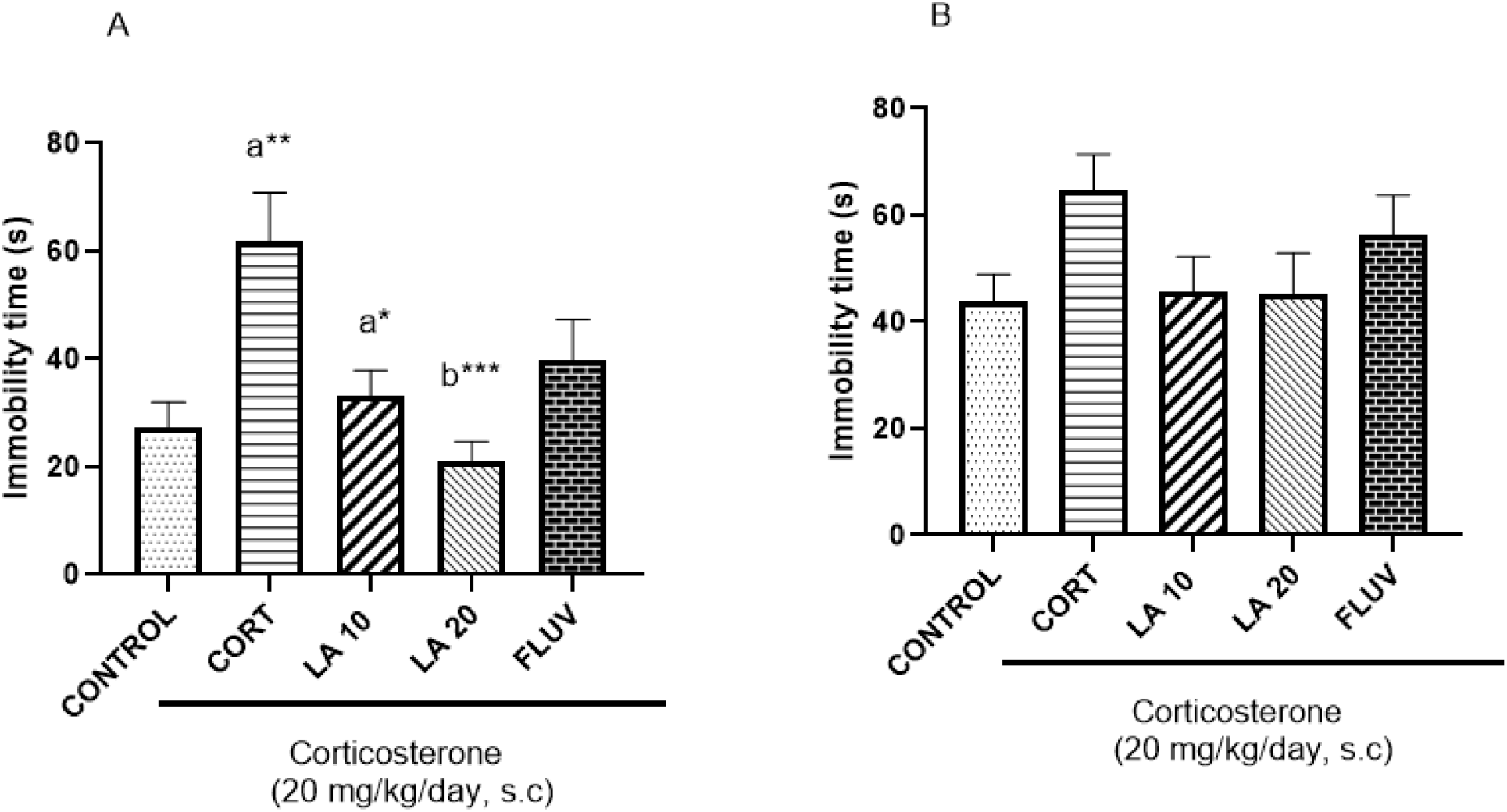
Effect of oral treatments with lauric acid (LA) and fluvoxamine on immobility time in the forced swimming test (2A) and tail suspension test (2B) in animals subjected to the corticosterone-induced depression model (20 mg/kg). Data are expressed as mean ± standard deviation and analyzed by one-way ANOVA followed by Tukey’s post hoc test (Fig. a), and as median ± interquartile range and analyzed using the Kruskal–Wallis test followed by Dunn’s test (Fig. b).The following letters were chosen to indicate the compared group ‘a’ show significant difference regarding to the control. ‘b’ show significant difference regarding to the group CORT. Significant values together the letters indicate: *p<0,01 e **p<0,001; ***p<0,0001.

The Tail Suspension Test (TST) is similar to the Forced Swimming Test (FST) and was developed as a methodological alternative to the FST in order to reduce bias related to the animals’ adaptation to swimming. In the present study, no significant alterations in mobility were observed in animals treated with lauric acid in the TST (Figure 2B).

### Open Field Test

The Open Field Test (OFT) is widely used to evaluate locomotor and exploratory activity in animals, allowing the exclusion of possible psychostimulant effects of the administered treatments. In this test, the parameters evaluated included horizontal exploratory activity (crossings), vertical exploration (rearing), self-cleaning behavior (grooming), and immobility (time during which the animals remained motionless). No significant alterations were observed in any of the parameters evaluated in the OFT (Figure 3).

**Fig 3:**
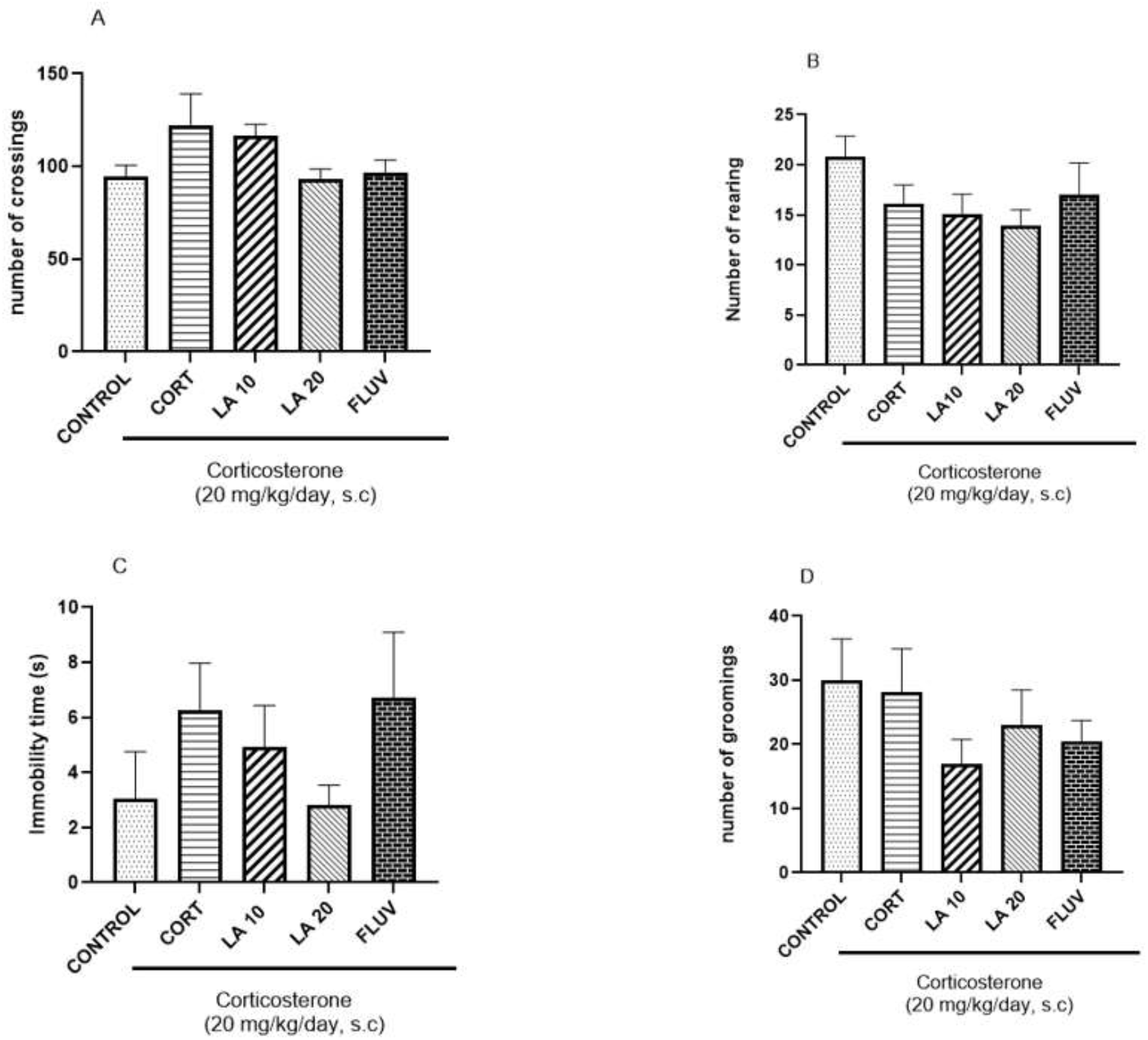
Number of crossings (a), rearings (b), immobility (c) and groomings (d) of animals in the open field test after chronic corticosterone injections CORT (20 mg/kg) and 9 days of oral administration of tested drugs LA: Lauric Acid (10 e 20 mg/kg) or FLUV: fluvoxamine (50 mg/kg). Data are expressed as mean ± standard deviation and analyzed by one-way ANOVA followed by Tukey’s post hoc test (Fig. b), and as median ± interquartile range and analyzed using the Kruskal–Wallis test followed by Dunn’s test (Fig. a,c,d).

### Neurochemical Assessments of Oxidative Stress

#### Effect of Lauric Acid on TBARS, Nitrite/Nitrate, and GSH Levels in Mouse Brains

The TBARS method was used to determine lipid peroxidation (cellular damage) through the measurement of malondialdehyde (MDA) concentration in brain regions such as the hippocampus, prefrontal cortex, and striatum. No significant alterations in MDA levels were observed in the hippocampus, prefrontal cortex, or striatum, as shown in figures 4A, 4B, and 4C, respectively. A significant alteration was observed only in the fluvoxamine-treated group when the prefrontal cortex was analyzed (Figure 4B).

**Fig 4:**
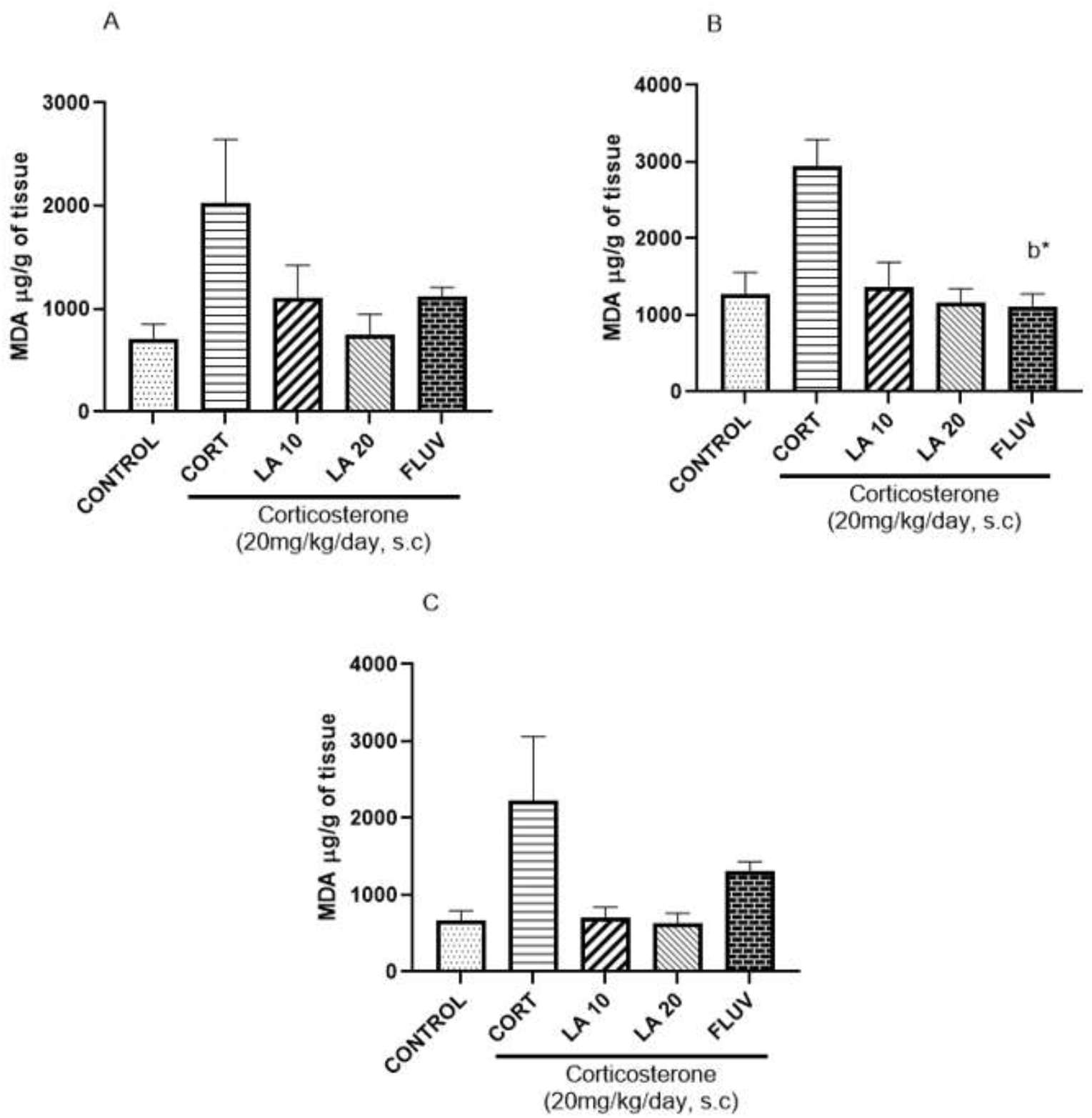
Effect of oral treatments with lauric acid (LA) and fluvoxamine in TBARS. Hipocampo (a),córtex pré-frontal (b), corpo estriado (c). Data are expressed as mean ± standard deviation and analyzed by one-way ANOVA followed by Tukey’s post hoc test (Fig. a,c), and as median ± interquartile range and analyzed using the Kruskal–Wallis test followed by Dunn’s test (Fig. b).The following letters were chosen to indicate the compared groups: ‘b’ indicates a significant difference compared to the CORT group. Significant values next to the letters indicate: * p< 0.01.

Nitrite/nitrate concentration in the brain regions was also evaluated as a measure of oxidative stress. Analysis of NO_2_^−^/NO_3_^−^ levels in the hippocampus and prefrontal cortex revealed no significant alterations (Figures 5A and 5B, respectively). However, a reduction in NO_2_^−^/NO_3_^−^ levels was observed in the striatum (Figure 5C). Likewise, no significant alterations in GSH levels were observed in the brain regions analyzed in this study, including the hippocampus (Figure 6A), prefrontal cortex (Figure 6B), and striatum (Figure 6C).

**Fig 5:**
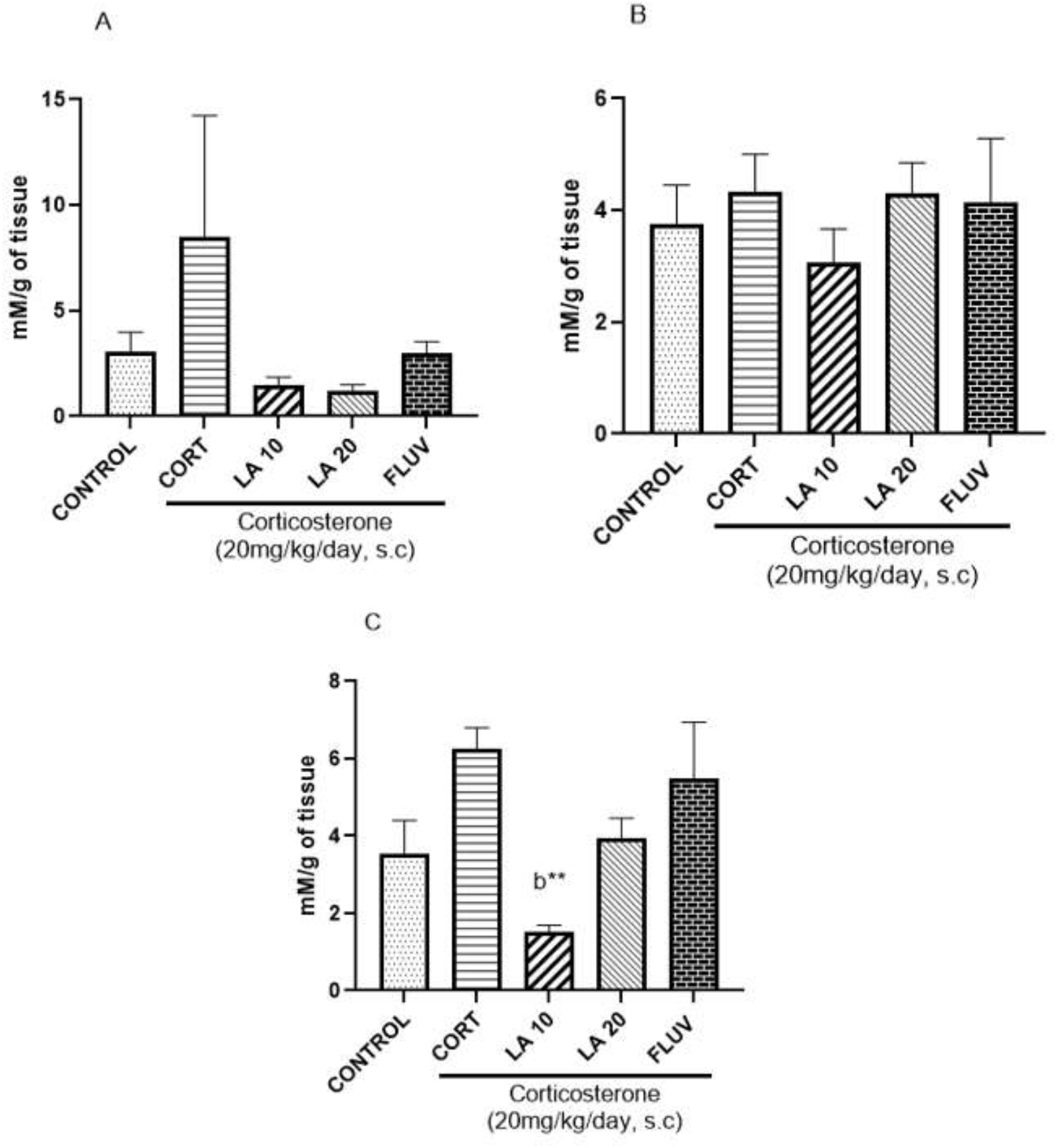
Effect of oral treatments with lauric acid (LA) and fluvoxamine in nitrite/nitrate. Hipocampo (a),córtex pré-frontal (b), corpo estriado (c). Data are expressed as mean ± standard deviation and analyzed by one-way ANOVA followed by Tukey’s post hoc test (Fig. a,c), and as median ± interquartile range and analyzed using the Kruskal–Wallis test followed by Dunn’s test (Fig. b).The following letters were chosen to indicate the compared groups: ‘b’ indicates a significant difference compared to the CORT group. Significant values next to the letters indicate: ** p< 0.001.

**Fig 6:**
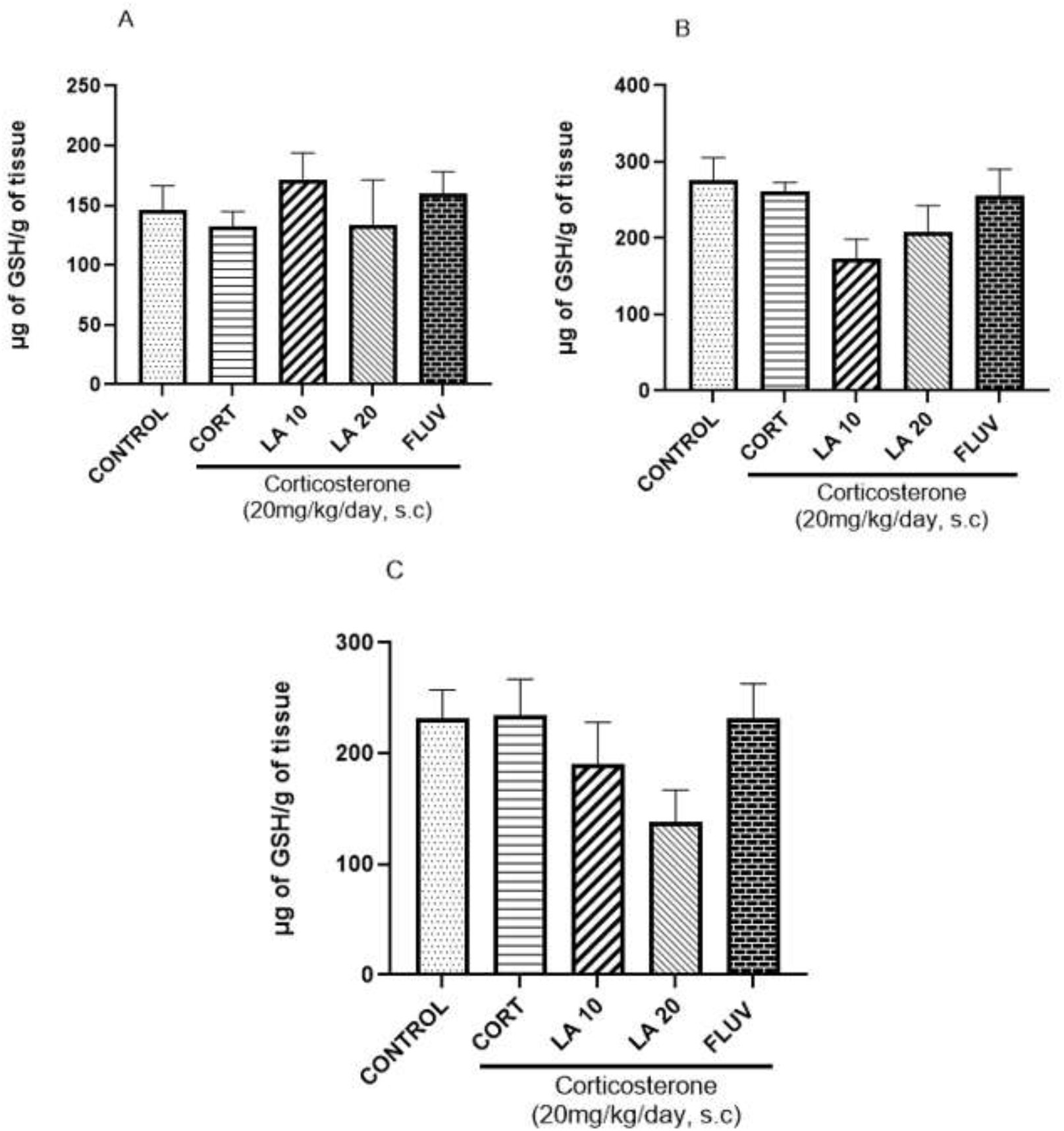
Effect of oral treatments with lauric acid (LA) and fluvoxamine in GSH. Hipocampo (a),córtex pré-frontal (b), corpo estriado (c). Data are expressed as mean ± standard deviation and analyzed by one-way ANOVA followed by Tukey’s post hoc test (Fig. b), and as median ± interquartile range and analyzed using the Kruskal–Wallis test followed by Dunn’s test (Fig. a,c).

## Discussion

In the present study, the effects of lauric acid (LA) were investigated in an experimental model of depressive-like behavior induced by exogenous corticosterone administration for 23 days, as described by Zhao et al. (2008).

Regarding the behavioral tests, both LA and fluvoxamine reduced the immobility time of the animals in the Forced Swimming Test (FST) compared to the CORT group. However, in the Tail Suspension Test (TST), no significant alterations in immobility behavior were observed.

The FST is widely used as a model of behavioral despair, in which animals are exposed to an unavoidable stressful situation by being placed in a water-filled cylinder without the possibility of escape, thereby inducing despair-like behavior (Can et al., 2012). In this context, both doses of LA reduced immobility time compared to the CORT group, showing an effect similar to that observed in the fluvoxamine-treated group, a selective serotonin reuptake inhibitor. These findings reinforce the potential antidepressant-like effect of LA. Furthermore, both the FST and TST are widely used tools for the preclinical screening of antidepressant compounds, as they provide relevant information regarding the adaptive capacity of animals when exposed to unavoidable stressful situations (Porsolt et al., 1977; Cryan et al., 2005).

In addition to evaluating antidepressant activity, it was necessary to investigate the locomotor activity of the animals in order to exclude the possibility of psychomotor interference resulting from treatment administration. Increased locomotor activity may be associated with psychostimulant effects, whereas reduced locomotion is often related to sedation (Bezerra et al., 2019). For this purpose, the Open Field Test (OFT) was employed, a widely used method for assessing exploratory activity, locomotion, and anxiety-like behavior (Kraeuter et al., 2019).

In the present study, LA did not promote significant alterations in the locomotor parameters evaluated in the OFT. However, Barreto (2020) demonstrated that LA administration at a dose of 10 mg/kg reduced the number of crossings in animals subjected to the OFT. This discrepancy may be related to the reduced number of animals per experimental group, a factor that may increase behavioral variability and compromise the statistical sensitivity of the analysis.

Regarding the number of crossings, LA at the dose of 20 mg/kg did not alter this parameter, suggesting that the antidepressant-like effect observed in this study is not associated with nonspecific psychostimulant effects. This finding reinforces the hypothesis that the reduction in depressive-like behavior promoted by LA occurs specifically, without interference in basal locomotor activity.

Another parameter evaluated in the OFT was the number of rearings, considered an indicator of vertical exploratory activity. This behavior corresponds to the number of times the animal remains supported exclusively on its hind limbs, without contact with the walls of the experimental apparatus (Sturman et al., 2017). It was observed that the CORT group reduced this parameter; however, treatment with LA at doses of 10 and 20 mg/kg was not able to reverse this alteration. Similarly, Barreto (2020) reported no alterations in the number of rearings following oral treatment with LA at the same doses. Thus, these results suggest that LA does not exert a significant effect on the vertical exploratory behavior of animals subjected to the experimental depression model.

Grooming behavior, related to self-cleaning activity, was also evaluated in the OFT. Alterations in this parameter have been associated with different behavioral manifestations observed in depression. Prolonged grooming episodes may be related to perseverative behaviors, whereas reduced grooming has been associated with anhedonia and decreased self-care, factors frequently linked to impaired social interactions (Kalueff et al., 2016; Cai et al., 2016). In the present study, no significant alterations were observed in the number of grooming episodes in the LA-treated groups. These results suggest that the treatment did not induce behavioral alterations related to anhedonia, particularly at the dose of 20 mg/kg, which exhibited a significant antidepressant-like effect.

In addition to behavioral alterations, neurochemical parameters related to oxidative and nitrosative stress were also investigated. Corticosterone administration increased lipid peroxidation levels, as expected, although it did not promote significant alterations in reduced glutathione (GSH) levels.

Regarding malondialdehyde (MDA) levels, no significant reduction was observed in the three evaluated brain regions — hippocampus, prefrontal cortex, and striatum — following treatment with LA at doses of 10 and 20 mg/kg, as well as in the fluvoxamine-treated group. These results contrast with the findings of Barreto (2020), who demonstrated reduced MDA levels in the striatum and prefrontal cortex following LA treatment. Similarly, Ribeiro (2024) observed reduced MDA levels in the hippocampus. Nevertheless, recent studies have demonstrated that LA exhibits antioxidant properties, reducing elevated MDA levels and restoring endogenous antioxidant systems such as superoxide dismutase (SOD) (Zaidi et al., 2020; Shaheryar et al., 2023).

In addition to reactive oxygen species, nitrite/nitrate (NO_2_^−^/NO_3_^−^) levels were also evaluated as indirect markers of nitrosative stress. Increased levels of these reactive species have been associated with cellular damage and the development of neuropsychiatric disorders, including depression (Moriyama et al., 2018). In the present study, the CORT group exhibited a significant increase in NO_2_^−^/NO_3_^−^ levels. Treatment with LA did not reduce these levels in the hippocampus or prefrontal cortex at either evaluated dose. However, in the striatum, a significant reduction was observed following LA administration at the dose of 10 mg/kg. Similar results were described by Barreto (2020), who demonstrated reduced nitrite/nitrate levels following preventive treatment with LA. In contrast, Ribeiro (2024) did not observe reductions in these markers in a preventive experimental model of depression. Additionally, Nishimura et al. (2018) demonstrated that LA was able to attenuate nitric oxide-induced damage resulting from microglial hyperactivation in an experimental model of Alzheimer’s disease. Taken together, these findings suggest that LA may exert modulatory effects on pathways related to nitrosative stress.

Reduced glutathione (GSH) levels were also evaluated in this study. No significant alterations in GSH levels were observed in the analyzed brain regions following LA treatment. Similar findings were reported by Ribeiro (2024), who demonstrated increased GSH levels only in the hippocampus following LA administration at the dose of 40 mg/kg, with no alterations observed in the prefrontal cortex or striatum. These findings suggest that higher doses of LA may be necessary to promote antioxidant effects mediated through the GSH pathway.

Taken together, the results obtained indicate that LA exerted a significant antidepressant-like effect in the experimental model used, without promoting nonspecific locomotor alterations. Furthermore, the observed neuroprotective effects appear to be partially related to modulation of nitrosative stress, although GSH-dependent antioxidant mechanisms were not identified under the experimental conditions evaluated. Therefore, additional studies are necessary to investigate other antioxidant pathways and molecular mechanisms potentially involved in the effects of LA on depressive disorders.

## Conclusion

The results presented in this study demonstrated that orally administered lauric acid (LA), at doses of 10 mg/kg and 20 mg/kg, has significant potential to reverse behavioral alterations induced by a corticosterone-induced model of depression over a short 23-day period in mice. Therefore, it can be concluded that LA exhibits antidepressant-like effects, as evidenced by the observed behavioral alterations. However, further studies are necessary to better understand the biochemical pathways involved in the mechanism of action of lauric acid.

