## Supplemental Data 1 for "Antidepressant Effects of Lauric Acid in a Corticosterone-Induced Murine Model of Depression: Behavioral and Neurochemical Insights": DADOS ESTATÍSTICOS.pdf

| Normality and Lognormality Tests | | A | B | C | D | E |
|---|---|---|---|---|---|---|
| | | CONTROL | CORT | LA 10 | LA 20 | FLUV |
| 1 | **Test for normal distribution** | | | | | |
| 2 | **Shapiro-Wilk test** | | | | | |
| 3 | W | 0.9411 | 0.9573 | 0.9578 | 0.9397 | 0.8990 |
| 4 | P value | 0.3025 | 0.5507 | 0.5599 | 0.2863 | 0.0553 |
| 5 | Passed normality test (alpha=0.05)? | Yes | Yes | Yes | Yes | Yes |
| 6 | P value summary | ns | ns | ns | ns | ns |
| 7 | | | | | | |
| 8 | **Kolmogorov-Smirnov test** | | | | | |
| 9 | KS distance | 0.1331 | 0.1343 | 0.1131 | 0.1237 | 0.2000 |
| 10 | P value | >0.1000 | >0.1000 | >0.1000 | >0.1000 | 0.0552 |
| 11 | Passed normality test (alpha=0.05)? | Yes | Yes | Yes | Yes | Yes |
| 12 | P value summary | ns | ns | ns | ns | ns |
| 13 | | | | | | |
| 14 | **Number of values** | 18 | 18 | 18 | 18 | 18 |

| Ordinary one-way ANOVA<br>ANOVA results | | | | | | |
|---|---|---|---|---|---|---|
| 1 | Table Analyzed | Nado Forçado | | | | |
| 2 | Data sets analyzed | A-E | | | | |
| 3 | | | | | | |
| 4 | **ANOVA summary** | | | | | |
| 5 | F | 6.339 | | | | |
| 6 | P value | 0.0002 | | | | |
| 7 | P value summary | *** | | | | |
| 8 | Significant diff. among means (P < 0.05)? | Yes | | | | |
| 9 | R squared | 0.2298 | | | | |
| 10 | | | | | | |
| 11 | **Brown-Forsythe test** | | | | | |
| 12 | F (DFn, DFd) | 4.959 (4, 85) | | | | |
| 13 | P value | 0.0012 | | | | |
| 14 | P value summary | ** | | | | |
| 15 | Are SDs significantly different (P < 0.05)? | Yes | | | | |
| 16 | | | | | | |
| 17 | **Bartlett's test** | | | | | |
| 18 | Bartlett's statistic (corrected) | 18.39 | | | | |
| 19 | P value | 0.0010 | | | | |
| 20 | P value summary | ** | | | | |
| 21 | Are SDs significantly different (P < 0.05)? | Yes | | | | |
| 22 | | | | | | |
| 23 | **ANOVA table** | **SS** | **DF** | **MS** | **F (DFn, DFd)** | **P value** |
| 24 | Treatment (between columns) | 17831 | 4 | 4458 | F (4, 85) = 6.339 | P=0.0002 |
| 25 | Residual (within columns) | 59777 | 85 | 703.3 | | |
| 26 | Total | 77608 | 89 | | | |
| 27 | | | | | | |
| 28 | **Data summary** | | | | | |
| 29 | Number of treatments (columns) | 5 | | | | |
| 30 | Number of values (total) | 90 | | | | |

| Ordinary one-way ANOVA<br>Multiple comparisons | | | | | |
|---|---|---|---|---|---|
| 1 | Number of families | 1 | | | |
| 2 | Number of comparisons per family | 10 | | | |
| 3 | Alpha | 0.05 | | | |
| 4 | | | | | |
| 5 | **Tukey's multiple comparisons test** | **Mean Diff.** | **95.00% CI of diff.** | **Significant?** | **Summary** |
| 6 | CONTROL vs. CORT | -34.61 | -59.25 to -9.973 | Yes | ** |
| 7 | CONTROL vs. LA10 | -5.944 | -30.58 to 18.69 | No | ns |
| 8 | CONTROL vs. LA20 | 6.278 | -18.36 to 30.92 | No | ns |
| 9 | CONTROL vs. FLUV | -12.39 | -37.03 to 12.25 | No | ns |
| 10 | CORT vs. LA10 | 28.67 | 4.029 to 53.30 | Yes | * |
| 11 | CORT vs. LA20 | 40.89 | 16.25 to 65.53 | Yes | *** |
| 12 | CORT vs. FLUV | 22.22 | -2.416 to 46.86 | No | ns |
| 13 | LA10 vs. LA20 | 12.22 | -12.42 to 36.86 | No | ns |
| 14 | LA10 vs. FLUV | -6.444 | -31.08 to 18.19 | No | ns |
| 15 | LA20 vs. FLUV | -18.67 | -43.30 to 5.971 | No | ns |
| 16 | | | | | |
| 17 | **Test details** | **Mean 1** | **Mean 2** | **Mean Diff.** | **SE of diff.** |
| 18 | CONTROL vs. CORT | 27.33 | 61.94 | -34.61 | 8.840 |
| 19 | CONTROL vs. LA10 | 27.33 | 33.28 | -5.944 | 8.840 |
| 20 | CONTROL vs. LA20 | 27.33 | 21.06 | 6.278 | 8.840 |
| 21 | CONTROL vs. FLUV | 27.33 | 39.72 | -12.39 | 8.840 |
| 22 | CORT vs. LA10 | 61.94 | 33.28 | 28.67 | 8.840 |
| 23 | CORT vs. LA20 | 61.94 | 21.06 | 40.89 | 8.840 |
| 24 | CORT vs. FLUV | 61.94 | 39.72 | 22.22 | 8.840 |
| 25 | LA10 vs. LA20 | 33.28 | 21.06 | 12.22 | 8.840 |
| 26 | LA10 vs. FLUV | 33.28 | 39.72 | -6.444 | 8.840 |
| 27 | LA20 vs. FLUV | 21.06 | 39.72 | -18.67 | 8.840 |

| | | | | |
|---|---|---|---|---|
| 1 | | | | |
| 2 | | | | |
| 3 | | | | |
| 4 | | | | |
| 5 | **Adjusted P Value** | | | |
| 6 | 0.0017 | A-B | | |
| 7 | 0.9618 | A-C | | |
| 8 | 0.9536 | A-D | | |
| 9 | 0.6284 | A-E | | |
| 10 | 0.0142 | B-C | | |
| 11 | 0.0001 | B-D | | |
| 12 | 0.0971 | B-E | | |
| 13 | 0.6404 | C-D | | |
| 14 | 0.9492 | C-E | | |
| 15 | 0.2247 | D-E | | |
| 16 | | | | |
| 17 | **n1** | **n2** | **q** | **DF** |
| 18 | 18 | 18 | 5.537 | 85 |
| 19 | 18 | 18 | 0.9510 | 85 |
| 20 | 18 | 18 | 1.004 | 85 |
| 21 | 18 | 18 | 1.982 | 85 |
| 22 | 18 | 18 | 4.586 | 85 |
| 23 | 18 | 18 | 6.542 | 85 |
| 24 | 18 | 18 | 3.555 | 85 |
| 25 | 18 | 18 | 1.955 | 85 |
| 26 | 18 | 18 | 1.031 | 85 |
| 27 | 18 | 18 | 2.986 | 85 |

| Normality and Lognormality Tests | | A<br>CONTROL | B<br>CORT | C<br>LA 10 | D<br>LA 20 | E<br>FLUV |
|---|---|---|---|---|---|---|
| 1 | **Test for normal distribution** | | | | | |
| 2 | **Anderson-Darling test** | | | | | |
| 3 | A2* | 0.4815 | 0.2884 | 0.7831 | 0.3170 | 0.2823 |
| 4 | P value | 0.2029 | 0.5756 | 0.0340 | 0.5108 | 0.5940 |
| 5 | Passed normality test (alpha=0.05)? | Yes | Yes | No | Yes | Yes |
| 6 | P value summary | ns | ns | * | ns | ns |
| 7 | | | | | | |
| 8 | **Shapiro-Wilk test** | | | | | |
| 9 | W | 0.9206 | 0.9692 | 0.8962 | 0.9477 | 0.9599 |
| 10 | P value | 0.1324 | 0.7831 | 0.0494 | 0.3898 | 0.5994 |
| 11 | Passed normality test (alpha=0.05)? | Yes | Yes | No | Yes | Yes |
| 12 | P value summary | ns | ns | * | ns | ns |
| 13 | | | | | | |
| 14 | **Number of values** | 18 | 18 | 18 | 18 | 18 |

| | Kruskal-Wallis test<br>ANOVA results | |
|---|---|---|
| 1 | Table Analyzed | Suspensão de cauda |
| 2 | | |
| 3 | **Kruskal-Wallis test** | |
| 4 | P value | 0.1791 |
| 5 | Exact or approximate P value? | Approximate |
| 6 | P value summary | ns |
| 7 | Do the medians vary signif. (P < 0.05)? | No |
| 8 | Number of groups | 5 |
| 9 | Kruskal-Wallis statistic | 6.281 |
| 10 | | |
| 11 | **Data summary** | |
| 12 | Number of treatments (columns) | 5 |
| 13 | Number of values (total) | 90 |

| **Kruskal-Wallis test**<br>Multiple comparisons | | | | | |
|---|---|---|---|---|---|
| 1 | Number of families | 1 | | | |
| 2 | Number of comparisons per family | 10 | | | |
| 3 | Alpha | 0.05 | | | |
| 4 | | | | | |
| 5 | **Dunn's multiple comparisons test** | **Mean rank diff.** | **Significant?** | **Summary** | **Adjusted P Value** |
| 6 | CONTROL vs. CORT | -17.81 | No | ns | 0.4082 |
| 7 | CONTROL vs. LA 10 | -1.167 | No | ns | >0.9999 |
| 8 | CONTROL vs. LA 20 | -0.4167 | No | ns | >0.9999 |
| 9 | CONTROL vs. FLUV | -8.944 | No | ns | >0.9999 |
| 10 | CORT vs. LA 10 | 16.64 | No | ns | 0.5596 |
| 11 | CORT vs. LA 20 | 17.39 | No | ns | 0.4577 |
| 12 | CORT vs. FLUV | 8.861 | No | ns | >0.9999 |
| 13 | LA 10 vs. LA 20 | 0.7500 | No | ns | >0.9999 |
| 14 | LA 10 vs. FLUV | -7.778 | No | ns | >0.9999 |
| 15 | LA 20 vs. FLUV | -8.528 | No | ns | >0.9999 |
| 16 | | | | | |
| 17 | **Test details** | **Mean rank 1** | **Mean rank 2** | **Mean rank diff.** | **n1** |
| 18 | CONTROL vs. CORT | 39.83 | 57.64 | -17.81 | 18 |
| 19 | CONTROL vs. LA 10 | 39.83 | 41.00 | -1.167 | 18 |
| 20 | CONTROL vs. LA 20 | 39.83 | 40.25 | -0.4167 | 18 |
| 21 | CONTROL vs. FLUV | 39.83 | 48.78 | -8.944 | 18 |
| 22 | CORT vs. LA 10 | 57.64 | 41.00 | 16.64 | 18 |
| 23 | CORT vs. LA 20 | 57.64 | 40.25 | 17.39 | 18 |
| 24 | CORT vs. FLUV | 57.64 | 48.78 | 8.861 | 18 |
| 25 | LA 10 vs. LA 20 | 41.00 | 40.25 | 0.7500 | 18 |
| 26 | LA 10 vs. FLUV | 41.00 | 48.78 | -7.778 | 18 |
| 27 | LA 20 vs. FLUV | 40.25 | 48.78 | -8.528 | 18 |

| | | |
|---|---|---|
| 1 | | |
| 2 | | |
| 3 | | |
| 4 | | |
| 5 | | |
| 6 | A-B | |
| 7 | A-C | |
| 8 | A-D | |
| 9 | A-E | |
| 10 | B-C | |
| 11 | B-D | |
| 12 | B-E | |
| 13 | C-D | |
| 14 | C-E | |
| 15 | D-E | |
| 16 | | |
| 17 | **n2** | **Z** |
| 18 | 18 | 2.045 |
| 19 | 18 | 0.1340 |
| 20 | 18 | 0.04786 |
| 21 | 18 | 1.027 |
| 22 | 18 | 1.911 |
| 23 | 18 | 1.997 |
| 24 | 18 | 1.018 |
| 25 | 18 | 0.08615 |
| 26 | 18 | 0.8934 |
| 27 | 18 | 0.9796 |

| Normality and Lognormality Tests | | A<br>CONTROL | B<br>CORT | C<br>LA10 | D<br>LA20 | E<br>FLUV |
|---|---|---|---|---|---|---|
| 1 | **Test for normal distribution** | | | | | |
| 2 | **Shapiro-Wilk test** | | | | | |
| 3 | W | 0.9486 | 0.9794 | 0.9475 | 0.9698 | 0.9198 |
| 4 | P value | 0.4036 | 0.9433 | 0.3870 | 0.7943 | 0.1283 |
| 5 | Passed normality test (alpha=0.05)? | Yes | Yes | Yes | Yes | Yes |
| 6 | P value summary | ns | ns | ns | ns | ns |
| 7 | | | | | | |
| 8 | **Number of values** | 18 | 18 | 18 | 18 | 18 |

| Ordinary one-way ANOVA<br>ANOVA results | | | | | | |
|---|---|---|---|---|---|---|
| 1 | Table Analyzed | Reanring | | | | |
| 2 | Data sets analyzed | A-E | | | | |
| 3 | | | | | | |
| 4 | **ANOVA summary** | | | | | |
| 5 | F | 1.455 | | | | |
| 6 | P value | 0.2230 | | | | |
| 7 | P value summary | ns | | | | |
| 8 | Significant diff. among means (P < 0.05)? | No | | | | |
| 9 | R squared | 0.06410 | | | | |
| 10 | | | | | | |
| 11 | **Brown-Forsythe test** | | | | | |
| 12 | F (DFn, DFd) | 2.758 (4, 85) | | | | |
| 13 | P value | 0.0329 | | | | |
| 14 | P value summary | * | | | | |
| 15 | Are SDs significantly different (P < 0.05)? | Yes | | | | |
| 16 | | | | | | |
| 17 | **Bartlett's test** | | | | | |
| 18 | Bartlett's statistic (corrected) | 9.945 | | | | |
| 19 | P value | 0.0414 | | | | |
| 20 | P value summary | * | | | | |
| 21 | Are SDs significantly different (P < 0.05)? | Yes | | | | |
| 22 | | | | | | |
| 23 | **ANOVA table** | **SS** | **DF** | **MS** | **F (DFn, DFd)** | **P value** |
| 24 | Treatment (between columns) | 499.6 | 4 | 124.9 | F (4, 85) = 1.455 | P=0.2230 |
| 25 | Residual (within columns) | 7294 | 85 | 85.81 | | |
| 26 | Total | 7793 | 89 | | | |
| 27 | | | | | | |
| 28 | **Data summary** | | | | | |
| 29 | Number of treatments (columns) | 5 | | | | |
| 30 | Number of values (total) | 90 | | | | |

| Ordinary one-way ANOVA<br>Multiple comparisons | | | | | |
|---|---|---|---|---|---|
| 1 | Number of families | 1 | | | |
| 2 | Number of comparisons per family | 10 | | | |
| 3 | Alpha | 0.05 | | | |
| 4 | | | | | |
| 5 | **Tukey's multiple comparisons test** | **Mean Diff.** | **95.00% CI of diff.** | **Significant?** | **Summary** |
| 6 | CONTROL vs. CORT | 4.667 | -3.940 to 13.27 | No | ns |
| 7 | CONTROL vs. LA10 | 5.778 | -2.828 to 14.38 | No | ns |
| 8 | CONTROL vs. LA20 | 6.889 | -1.717 to 15.50 | No | ns |
| 9 | CONTROL vs. FLUV | 3.778 | -4.828 to 12.38 | No | ns |
| 10 | CORT vs. LA10 | 1.111 | -7.495 to 9.717 | No | ns |
| 11 | CORT vs. LA20 | 2.222 | -6.384 to 10.83 | No | ns |
| 12 | CORT vs. FLUV | -0.8889 | -9.495 to 7.717 | No | ns |
| 13 | LA10 vs. LA20 | 1.111 | -7.495 to 9.717 | No | ns |
| 14 | LA10 vs. FLUV | -2.000 | -10.61 to 6.606 | No | ns |
| 15 | LA20 vs. FLUV | -3.111 | -11.72 to 5.495 | No | ns |
| 16 | | | | | |
| 17 | **Test details** | **Mean 1** | **Mean 2** | **Mean Diff.** | **SE of diff.** |
| 18 | CONTROL vs. CORT | 20.83 | 16.17 | 4.667 | 3.088 |
| 19 | CONTROL vs. LA10 | 20.83 | 15.06 | 5.778 | 3.088 |
| 20 | CONTROL vs. LA20 | 20.83 | 13.94 | 6.889 | 3.088 |
| 21 | CONTROL vs. FLUV | 20.83 | 17.06 | 3.778 | 3.088 |
| 22 | CORT vs. LA10 | 16.17 | 15.06 | 1.111 | 3.088 |
| 23 | CORT vs. LA20 | 16.17 | 13.94 | 2.222 | 3.088 |
| 24 | CORT vs. FLUV | 16.17 | 17.06 | -0.8889 | 3.088 |
| 25 | LA10 vs. LA20 | 15.06 | 13.94 | 1.111 | 3.088 |
| 26 | LA10 vs. FLUV | 15.06 | 17.06 | -2.000 | 3.088 |
| 27 | LA20 vs. FLUV | 13.94 | 17.06 | -3.111 | 3.088 |

| | | | | |
|---|---|---|---|---|
| 1 | | | | |
| 2 | | | | |
| 3 | | | | |
| 4 | | | | |
| 5 | **Adjusted P Value** | | | |
| 6 | 0.5581 | A-B | | |
| 7 | 0.3407 | A-C | | |
| 8 | 0.1784 | A-D | | |
| 9 | 0.7378 | A-E | | |
| 10 | 0.9963 | B-C | | |
| 11 | 0.9514 | B-D | | |
| 12 | 0.9985 | B-E | | |
| 13 | 0.9963 | C-D | | |
| 14 | 0.9666 | C-E | | |
| 15 | 0.8512 | D-E | | |
| 16 | | | | |
| 17 | **n1** | **n2** | **q** | **DF** |
| 18 | 18 | 18 | 2.137 | 85 |
| 19 | 18 | 18 | 2.646 | 85 |
| 20 | 18 | 18 | 3.155 | 85 |
| 21 | 18 | 18 | 1.730 | 85 |
| 22 | 18 | 18 | 0.5089 | 85 |
| 23 | 18 | 18 | 1.018 | 85 |
| 24 | 18 | 18 | 0.4071 | 85 |
| 25 | 18 | 18 | 0.5089 | 85 |
| 26 | 18 | 18 | 0.9160 | 85 |
| 27 | 18 | 18 | 1.425 | 85 |

| Normality and Lognormality Tests | | A | B | C | D | E |
|---|---|---|---|---|---|---|
| | | CONTROL | CORT | LA 10 | LA 20 | FLUV |
| 1 | **Test for normal distribution** | | | | | |
| 2 | **Shapiro-Wilk test** | | | | | |
| 3 | W | 0.9376 | 0.7348 | 0.9238 | 0.9227 | 0.9679 |
| 4 | P value | 0.2639 | 0.0002 | 0.1506 | 0.1443 | 0.7582 |
| 5 | Passed normality test (alpha=0.05)? | Yes | No | Yes | Yes | Yes |
| 6 | P value summary | ns | *** | ns | ns | ns |
| 7 | | | | | | |
| 8 | **Kolmogorov-Smirnov test** | | | | | |
| 9 | KS distance | 0.1154 | 0.2858 | 0.1766 | 0.1656 | 0.1101 |
| 10 | P value | >0.1000 | 0.0004 | >0.1000 | >0.1000 | >0.1000 |
| 11 | Passed normality test (alpha=0.05)? | Yes | No | Yes | Yes | Yes |
| 12 | P value summary | ns | *** | ns | ns | ns |
| 13 | | | | | | |
| 14 | **Number of values** | 18 | 18 | 18 | 18 | 18 |

**Kruskal-Wallis test**
ANOVA results

| | | |
|---|---|---|
| 1 | Table Analyzed | Crossings |
| 2 | | |
| 3 | **Kruskal-Wallis test** | |
| 4 | P value | 0.0678 |
| 5 | Exact or approximate P value? | Approximate |
| 6 | P value summary | ns |
| 7 | Do the medians vary signif. (P < 0.05)? | No |
| 8 | Number of groups | 5 |
| 9 | Kruskal-Wallis statistic | 8.746 |
| 10 | | |
| 11 | **Data summary** | |
| 12 | Number of treatments (columns) | 5 |
| 13 | Number of values (total) | 90 |

| **Kruskal-Wallis test**<br>Multiple comparisons | | | | | |
|---|---|---|---|---|---|
| 1 | Number of families | 1 | | | |
| 2 | Number of comparisons per family | 10 | | | |
| 3 | Alpha | 0.05 | | | |
| 4 | | | | | |
| 5 | **Dunn's multiple comparisons test** | **Mean rank diff.** | **Significant?** | **Summary** | **Adjusted P Value** |
| 6 | CONTROL vs. CORT | -9.139 | No | ns | >0.9999 |
| 7 | CONTROL vs. LA 10 | -18.72 | No | ns | 0.3150 |
| 8 | CONTROL vs. LA 20 | 4.806 | No | ns | >0.9999 |
| 9 | CONTROL vs. FLUV | -2.778 | No | ns | >0.9999 |
| 10 | CORT vs. LA 10 | -9.583 | No | ns | >0.9999 |
| 11 | CORT vs. LA 20 | 13.94 | No | ns | >0.9999 |
| 12 | CORT vs. FLUV | 6.361 | No | ns | >0.9999 |
| 13 | LA 10 vs. LA 20 | 23.53 | No | ns | 0.0688 |
| 14 | LA 10 vs. FLUV | 15.94 | No | ns | 0.6701 |
| 15 | LA 20 vs. FLUV | -7.583 | No | ns | >0.9999 |
| 16 | | | | | |
| 17 | **Test details** | **Mean rank 1** | **Mean rank 2** | **Mean rank diff.** | **n1** |
| 18 | CONTROL vs. CORT | 40.33 | 49.47 | -9.139 | 18 |
| 19 | CONTROL vs. LA 10 | 40.33 | 59.06 | -18.72 | 18 |
| 20 | CONTROL vs. LA 20 | 40.33 | 35.53 | 4.806 | 18 |
| 21 | CONTROL vs. FLUV | 40.33 | 43.11 | -2.778 | 18 |
| 22 | CORT vs. LA 10 | 49.47 | 59.06 | -9.583 | 18 |
| 23 | CORT vs. LA 20 | 49.47 | 35.53 | 13.94 | 18 |
| 24 | CORT vs. FLUV | 49.47 | 43.11 | 6.361 | 18 |
| 25 | LA 10 vs. LA 20 | 59.06 | 35.53 | 23.53 | 18 |
| 26 | LA 10 vs. FLUV | 59.06 | 43.11 | 15.94 | 18 |
| 27 | LA 20 vs. FLUV | 35.53 | 43.11 | -7.583 | 18 |

| | | |
|---|---|---|
| 1 | | |
| 2 | | |
| 3 | | |
| 4 | | |
| 5 | | |
| 6 | A-B | |
| 7 | A-C | |
| 8 | A-D | |
| 9 | A-E | |
| 10 | B-C | |
| 11 | B-D | |
| 12 | B-E | |
| 13 | C-D | |
| 14 | C-E | |
| 15 | D-E | |
| 16 | | |
| 17 | **n2** | **Z** |
| 18 | 18 | 1.050 |
| 19 | 18 | 2.151 |
| 20 | 18 | 0.5520 |
| 21 | 18 | 0.3191 |
| 22 | 18 | 1.101 |
| 23 | 18 | 1.602 |
| 24 | 18 | 0.7307 |
| 25 | 18 | 2.703 |
| 26 | 18 | 1.832 |
| 27 | 18 | 0.8711 |

| Normality and Lognormality Tests | | A | B | C | D | E |
|---|---|---|---|---|---|---|
| | | CONTROL | CORT | LA 10 | LA 20 | FLUV |
| 1 | **Test for normal distribution** | | | | | |
| 2 | **Shapiro-Wilk test** | | | | | |
| 3 | W | 0.4469 | 0.8224 | 0.7717 | 0.8323 | 0.6533 |
| 4 | P value | <0.0001 | 0.0032 | 0.0006 | 0.0045 | <0.0001 |
| 5 | Passed normality test (alpha=0.05)? | No | No | No | No | No |
| 6 | P value summary | **** | ** | *** | ** | **** |
| 7 | | | | | | |
| 8 | **Number of values** | 18 | 18 | 18 | 18 | 18 |

| Kruskal-Wallis test<br>ANOVA results | | |
|---|---|---|
| 1 | Table Analyzed | Immobility time |
| 2 | | |
| 3 | **Kruskal-Wallis test** | |
| 4 | P value | 0.1925 |
| 5 | Exact or approximate P value? | Approximate |
| 6 | P value summary | ns |
| 7 | Do the medians vary signif. (P < 0.05)? | No |
| 8 | Number of groups | 5 |
| 9 | Kruskal-Wallis statistic | 6.091 |
| 10 | | |
| 11 | **Data summary** | |
| 12 | Number of treatments (columns) | 5 |
| 13 | Number of values (total) | 90 |

| Kruskal-Wallis test<br>Multiple comparisons | | | | | |
|---|---|---|---|---|---|
| 1 | Number of families | 1 | | | |
| 2 | Number of comparisons per family | 10 | | | |
| 3 | Alpha | 0.05 | | | |
| 4 | | | | | |
| 5 | **Dunn's multiple comparisons test** | **Mean rank diff.** | **Significant?** | **Summary** | **Adjusted P Value** |
| 6 | CONTROL vs. CORT | -18.78 | No | ns | 0.2768 |
| 7 | CONTROL vs. LA 10 | -13.39 | No | ns | >0.9999 |
| 8 | CONTROL vs. LA 20 | -6.361 | No | ns | >0.9999 |
| 9 | CONTROL vs. FLUV | -14.67 | No | ns | 0.8548 |
| 10 | CORT vs. LA 10 | 5.389 | No | ns | >0.9999 |
| 11 | CORT vs. LA 20 | 12.42 | No | ns | >0.9999 |
| 12 | CORT vs. FLUV | 4.111 | No | ns | >0.9999 |
| 13 | LA 10 vs. LA 20 | 7.028 | No | ns | >0.9999 |
| 14 | LA 10 vs. FLUV | -1.278 | No | ns | >0.9999 |
| 15 | LA 20 vs. FLUV | -8.306 | No | ns | >0.9999 |
| 16 | | | | | |
| 17 | **Test details** | **Mean rank 1** | **Mean rank 2** | **Mean rank diff.** | **n1** |
| 18 | CONTROL vs. CORT | 34.86 | 53.64 | -18.78 | 18 |
| 19 | CONTROL vs. LA 10 | 34.86 | 48.25 | -13.39 | 18 |
| 20 | CONTROL vs. LA 20 | 34.86 | 41.22 | -6.361 | 18 |
| 21 | CONTROL vs. FLUV | 34.86 | 49.53 | -14.67 | 18 |
| 22 | CORT vs. LA 10 | 53.64 | 48.25 | 5.389 | 18 |
| 23 | CORT vs. LA 20 | 53.64 | 41.22 | 12.42 | 18 |
| 24 | CORT vs. FLUV | 53.64 | 49.53 | 4.111 | 18 |
| 25 | LA 10 vs. LA 20 | 48.25 | 41.22 | 7.028 | 18 |
| 26 | LA 10 vs. FLUV | 48.25 | 49.53 | -1.278 | 18 |
| 27 | LA 20 vs. FLUV | 41.22 | 49.53 | -8.306 | 18 |

| | | |
|---|---|---|
| 1 | | |
| 2 | | |
| 3 | | |
| 4 | | |
| 5 | | |
| 6 | A-B | |
| 7 | A-C | |
| 8 | A-D | |
| 9 | A-E | |
| 10 | B-C | |
| 11 | B-D | |
| 12 | B-E | |
| 13 | C-D | |
| 14 | C-E | |
| 15 | D-E | |
| 16 | | |
| 17 | **n2** | **Z** |
| 18 | 18 | 2.202 |
| 19 | 18 | 1.570 |
| 20 | 18 | 0.7459 |
| 21 | 18 | 1.720 |
| 22 | 18 | 0.6319 |
| 23 | 18 | 1.456 |
| 24 | 18 | 0.4820 |
| 25 | 18 | 0.8240 |
| 26 | 18 | 0.1498 |
| 27 | 18 | 0.9739 |

| Normality and Lognormality Tests | | A | B | C | D | E |
|---|---|---|---|---|---|---|
| | | CONTROL | CORT | LA 10 | LA 20 | FLUV |
| 1 | **Test for normal distribution** | | | | | |
| 2 | **Shapiro-Wilk test** | | | | | |
| 3 | W | 0.6991 | 0.7536 | 0.7089 | 0.7398 | 0.9267 |
| 4 | P value | <0.0001 | 0.0004 | 0.0001 | 0.0002 | 0.1698 |
| 5 | Passed normality test (alpha=0.05)? | No | No | No | No | Yes |
| 6 | P value summary | **** | *** | *** | *** | ns |
| 7 | | | | | | |
| 8 | **Number of values** | 18 | 18 | 18 | 18 | 18 |

| Kruskal-Wallis test<br>ANOVA results | | |
|---|---|---|
| 1 | Table Analyzed | Groomings |
| 2 | | |
| 3 | **Kruskal-Wallis test** | |
| 4 | P value | 0.2088 |
| 5 | Exact or approximate P value? | Approximate |
| 6 | P value summary | ns |
| 7 | Do the medians vary signif. (P < 0.05)? | No |
| 8 | Number of groups | 5 |
| 9 | Kruskal-Wallis statistic | 5.874 |
| 10 | | |
| 11 | **Data summary** | |
| 12 | Number of treatments (columns) | 5 |
| 13 | Number of values (total) | 90 |

| **Kruskal-Wallis test**<br>Multiple comparisons | | | | | |
|---|---|---|---|---|---|
| 1 | Number of families | 1 | | | |
| 2 | Number of comparisons per family | 10 | | | |
| 3 | Alpha | 0.05 | | | |
| 4 | | | | | |
| 5 | **Dunn's multiple comparisons test** | **Mean rank diff.** | **Significant?** | **Summary** | **Adjusted P Value** |
| 6 | CONTROL vs. CORT | 7.111 | No | ns | >0.9999 |
| 7 | CONTROL vs. LA 10 | 20.31 | No | ns | 0.1963 |
| 8 | CONTROL vs. LA 20 | 12.42 | No | ns | >0.9999 |
| 9 | CONTROL vs. FLUV | 11.56 | No | ns | >0.9999 |
| 10 | CORT vs. LA 10 | 13.19 | No | ns | >0.9999 |
| 11 | CORT vs. LA 20 | 5.306 | No | ns | >0.9999 |
| 12 | CORT vs. FLUV | 4.444 | No | ns | >0.9999 |
| 13 | LA 10 vs. LA 20 | -7.889 | No | ns | >0.9999 |
| 14 | LA 10 vs. FLUV | -8.750 | No | ns | >0.9999 |
| 15 | LA 20 vs. FLUV | -0.8611 | No | ns | >0.9999 |
| 16 | | | | | |
| 17 | **Test details** | **Mean rank 1** | **Mean rank 2** | **Mean rank diff.** | **n1** |
| 18 | CONTROL vs. CORT | 55.78 | 48.67 | 7.111 | 18 |
| 19 | CONTROL vs. LA 10 | 55.78 | 35.47 | 20.31 | 18 |
| 20 | CONTROL vs. LA 20 | 55.78 | 43.36 | 12.42 | 18 |
| 21 | CONTROL vs. FLUV | 55.78 | 44.22 | 11.56 | 18 |
| 22 | CORT vs. LA 10 | 48.67 | 35.47 | 13.19 | 18 |
| 23 | CORT vs. LA 20 | 48.67 | 43.36 | 5.306 | 18 |
| 24 | CORT vs. FLUV | 48.67 | 44.22 | 4.444 | 18 |
| 25 | LA 10 vs. LA 20 | 35.47 | 43.36 | -7.889 | 18 |
| 26 | LA 10 vs. FLUV | 35.47 | 44.22 | -8.750 | 18 |
| 27 | LA 20 vs. FLUV | 43.36 | 44.22 | -0.8611 | 18 |

| | | |
|---|---|---|
| 1 | | |
| 2 | | |
| 3 | | |
| 4 | | |
| 5 | | |
| 6 | A-B | |
| 7 | A-C | |
| 8 | A-D | |
| 9 | A-E | |
| 10 | B-C | |
| 11 | B-D | |
| 12 | B-E | |
| 13 | C-D | |
| 14 | C-E | |
| 15 | D-E | |
| 16 | | |
| 17 | **n2** | **Z** |
| 18 | 18 | 0.8171 |
| 19 | 18 | 2.333 |
| 20 | 18 | 1.427 |
| 21 | 18 | 1.328 |
| 22 | 18 | 1.516 |
| 23 | 18 | 0.6096 |
| 24 | 18 | 0.5107 |
| 25 | 18 | 0.9065 |
| 26 | 18 | 1.005 |
| 27 | 18 | 0.09895 |

| Normality and Lognormality Tests | | A | B | C | D | E |
|---|---|---|---|---|---|---|
| | | CONTROL | CORT | LA 10 | LA 20 | FLUV |
| 1 | **Test for normal distribution** | | | | | |
| 2 | **Shapiro-Wilk test** | | | | | |
| 3 | W | 0.9209 | 0.9164 | 0.8405 | 0.9118 | 0.9611 |
| 4 | P value | 0.5120 | 0.4801 | 0.1317 | 0.4485 | 0.8283 |
| 5 | Passed normality test (alpha=0.05)? | Yes | Yes | Yes | Yes | Yes |
| 6 | P value summary | ns | ns | ns | ns | ns |
| 7 | | | | | | |
| 8 | **Number of values** | 6 | 6 | 6 | 6 | 6 |

| Ordinary one-way ANOVA<br>ANOVA results | | | | | | |
|---|---|---|---|---|---|---|
| 1 | Table Analyzed | TBARS HIPOCAMP | | | | |
| 2 | Data sets analyzed | A-E | | | | |
| 3 | | | | | | |
| 4 | **ANOVA summary** | | | | | |
| 5 | F | 2.507 | | | | |
| 6 | P value | 0.0676 | | | | |
| 7 | P value summary | ns | | | | |
| 8 | Significant diff. among means (P < 0.05)? | No | | | | |
| 9 | R squared | 0.2863 | | | | |
| 10 | | | | | | |
| 11 | **Brown-Forsythe test** | | | | | |
| 12 | F (DFn, DFd) | 3.777 (4, 25) | | | | |
| 13 | P value | 0.0155 | | | | |
| 14 | P value summary | * | | | | |
| 15 | Are SDs significantly different (P < 0.05)? | Yes | | | | |
| 16 | | | | | | |
| 17 | **Bartlett's test** | | | | | |
| 18 | Bartlett's statistic (corrected) | 19.37 | | | | |
| 19 | P value | 0.0007 | | | | |
| 20 | P value summary | *** | | | | |
| 21 | Are SDs significantly different (P < 0.05)? | Yes | | | | |
| 22 | | | | | | |
| 23 | **ANOVA table** | **SS** | **DF** | **MS** | **F (DFn, DFd)** | **P value** |
| 24 | Treatment (between columns) | 6737951 | 4 | 1684488 | F (4, 25) = 2.507 | P=0.0676 |
| 25 | Residual (within columns) | 16799452 | 25 | 671978 | | |
| 26 | Total | 23537403 | 29 | | | |
| 27 | | | | | | |
| 28 | **Data summary** | | | | | |
| 29 | Number of treatments (columns) | 5 | | | | |
| 30 | Number of values (total) | 30 | | | | |

| Ordinary one-way ANOVA<br>Multiple comparisons | | | | | |
|---|---|---|---|---|---|
| 1 | Number of families | 1 | | | |
| 2 | Number of comparisons per family | 10 | | | |
| 3 | Alpha | 0.05 | | | |
| 4 | | | | | |
| 5 | **Tukey's multiple comparisons test** | **Mean Diff.** | **95.00% CI of diff.** | **Significant?** | **Summary** |
| 6 | CONTROL vs. CORT | -1321 | -2711 to 68.79 | No | ns |
| 7 | CONTROL vs. LA 10 | -397.8 | -1788 to 992.1 | No | ns |
| 8 | CONTROL vs. LA 20 | -51.83 | -1442 to 1338 | No | ns |
| 9 | CONTROL vs. FLUV | -417.3 | -1807 to 972.6 | No | ns |
| 10 | CORT vs. LA 10 | 923.3 | -466.6 to 2313 | No | ns |
| 11 | CORT vs. LA 20 | 1269 | -120.6 to 2659 | No | ns |
| 12 | CORT vs. FLUV | 903.8 | -486.1 to 2294 | No | ns |
| 13 | LA 10 vs. LA 20 | 346.0 | -1044 to 1736 | No | ns |
| 14 | LA 10 vs. FLUV | -19.50 | -1409 to 1370 | No | ns |
| 15 | LA 20 vs. FLUV | -365.5 | -1755 to 1024 | No | ns |
| 16 | | | | | |
| 17 | **Test details** | **Mean 1** | **Mean 2** | **Mean Diff.** | **SE of diff.** |
| 18 | CONTROL vs. CORT | 703.3 | 2025 | -1321 | 473.3 |
| 19 | CONTROL vs. LA 10 | 703.3 | 1101 | -397.8 | 473.3 |
| 20 | CONTROL vs. LA 20 | 703.3 | 755.2 | -51.83 | 473.3 |
| 21 | CONTROL vs. FLUV | 703.3 | 1121 | -417.3 | 473.3 |
| 22 | CORT vs. LA 10 | 2025 | 1101 | 923.3 | 473.3 |
| 23 | CORT vs. LA 20 | 2025 | 755.2 | 1269 | 473.3 |
| 24 | CORT vs. FLUV | 2025 | 1121 | 903.8 | 473.3 |
| 25 | LA 10 vs. LA 20 | 1101 | 755.2 | 346.0 | 473.3 |
| 26 | LA 10 vs. FLUV | 1101 | 1121 | -19.50 | 473.3 |
| 27 | LA 20 vs. FLUV | 755.2 | 1121 | -365.5 | 473.3 |

| | | | | |
|---|---|---|---|---|
| 1 | | | | |
| 2 | | | | |
| 3 | | | | |
| 4 | | | | |
| 5 | **Adjusted P Value** | | | |
| 6 | 0.0682 | A-B | | |
| 7 | 0.9153 | A-C | | |
| 8 | >0.9999 | A-D | | |
| 9 | 0.9009 | A-E | | |
| 10 | 0.3181 | B-C | | |
| 11 | 0.0855 | B-D | | |
| 12 | 0.3383 | B-E | | |
| 13 | 0.9472 | C-D | | |
| 14 | >0.9999 | C-E | | |
| 15 | 0.9362 | D-E | | |
| 16 | | | | |
| 17 | **n1** | **n2** | **q** | **DF** |
| 18 | 6 | 6 | 3.948 | 25 |
| 19 | 6 | 6 | 1.189 | 25 |
| 20 | 6 | 6 | 0.1549 | 25 |
| 21 | 6 | 6 | 1.247 | 25 |
| 22 | 6 | 6 | 2.759 | 25 |
| 23 | 6 | 6 | 3.793 | 25 |
| 24 | 6 | 6 | 2.701 | 25 |
| 25 | 6 | 6 | 1.034 | 25 |
| 26 | 6 | 6 | 0.05827 | 25 |
| 27 | 6 | 6 | 1.092 | 25 |

| Normality and Lognormality Tests | | A | B | C | D | E |
|---|---|---|---|---|---|---|
| | | CONTROL | CORT | LA 10 | LA 20 | FLUV |
| 1 | **Test for normal distribution** | | | | | |
| 2 | **Shapiro-Wilk test** | | | | | |
| 3 | W | 0.7883 | 0.7995 | 0.8839 | 0.9367 | 0.8635 |
| 4 | P value | 0.0460 | 0.0582 | 0.2873 | 0.6330 | 0.2017 |
| 5 | Passed normality test (alpha=0.05)? | No | Yes | Yes | Yes | Yes |
| 6 | P value summary | * | ns | ns | ns | ns |
| 7 | | | | | | |
| 8 | **Number of values** | 6 | 6 | 6 | 6 | 6 |

| | **Kruskal-Wallis test**<br>ANOVA results | |
|---|---|---|
| 1 | Table Analyzed | TBARS CÓRTEX PRÉ - FRONTA |
| 2 | | |
| 3 | **Kruskal-Wallis test** | |
| 4 | P value | 0.0166 |
| 5 | Exact or approximate P value? | Approximate |
| 6 | P value summary | * |
| 7 | Do the medians vary signif. (P < 0.05)? | Yes |
| 8 | Number of groups | 5 |
| 9 | Kruskal-Wallis statistic | 12.11 |
| 10 | | |
| 11 | **Data summary** | |
| 12 | Number of treatments (columns) | 5 |
| 13 | Number of values (total) | 30 |

| Kruskal-Wallis test<br>Multiple comparisons | | | | | |
|---|---|---|---|---|---|
| 1 | Number of families | 1 | | | |
| 2 | Number of comparisons per family | 10 | | | |
| 3 | Alpha | 0.05 | | | |
| 4 | | | | | |
| 5 | **Dunn's multiple comparisons test** | **Mean rank diff.** | **Significant?** | **Summary** | **Adjusted P Value** |
| 6 | CONTROL vs. CORT | -13.50 | No | ns | 0.0791 |
| 7 | CONTROL vs. LA 10 | -1.333 | No | ns | >0.9999 |
| 8 | CONTROL vs. LA 20 | 0.5000 | No | ns | >0.9999 |
| 9 | CONTROL vs. FLUV | 1.833 | No | ns | >0.9999 |
| 10 | CORT vs. LA 10 | 12.17 | No | ns | 0.1668 |
| 11 | CORT vs. LA 20 | 14.00 | No | ns | 0.0588 |
| 12 | CORT vs. FLUV | 15.33 | Yes | * | 0.0255 |
| 13 | LA 10 vs. LA 20 | 1.833 | No | ns | >0.9999 |
| 14 | LA 10 vs. FLUV | 3.167 | No | ns | >0.9999 |
| 15 | LA 20 vs. FLUV | 1.333 | No | ns | >0.9999 |
| 16 | | | | | |
| 17 | **Test details** | **Mean rank 1** | **Mean rank 2** | **Mean rank diff.** | **n1** |
| 18 | CONTROL vs. CORT | 13.00 | 26.50 | -13.50 | 6 |
| 19 | CONTROL vs. LA 10 | 13.00 | 14.33 | -1.333 | 6 |
| 20 | CONTROL vs. LA 20 | 13.00 | 12.50 | 0.5000 | 6 |
| 21 | CONTROL vs. FLUV | 13.00 | 11.17 | 1.833 | 6 |
| 22 | CORT vs. LA 10 | 26.50 | 14.33 | 12.17 | 6 |
| 23 | CORT vs. LA 20 | 26.50 | 12.50 | 14.00 | 6 |
| 24 | CORT vs. FLUV | 26.50 | 11.17 | 15.33 | 6 |
| 25 | LA 10 vs. LA 20 | 14.33 | 12.50 | 1.833 | 6 |
| 26 | LA 10 vs. FLUV | 14.33 | 11.17 | 3.167 | 6 |
| 27 | LA 20 vs. FLUV | 12.50 | 11.17 | 1.333 | 6 |

| | | |
|---|---|---|
| 1 | | |
| 2 | | |
| 3 | | |
| 4 | | |
| 5 | | |
| 6 | A-B | |
| 7 | A-C | |
| 8 | A-D | |
| 9 | A-E | |
| 10 | B-C | |
| 11 | B-D | |
| 12 | B-E | |
| 13 | C-D | |
| 14 | C-E | |
| 15 | D-E | |
| 16 | | |
| 17 | **n2** | **Z** |
| 18 | 6 | 2.656 |
| 19 | 6 | 0.2623 |
| 20 | 6 | 0.09837 |
| 21 | 6 | 0.3607 |
| 22 | 6 | 2.394 |
| 23 | 6 | 2.754 |
| 24 | 6 | 3.017 |
| 25 | 6 | 0.3607 |
| 26 | 6 | 0.6230 |
| 27 | 6 | 0.2623 |

| Normality and Lognormality Tests | | A | B | C | D | E |
|---|---|---|---|---|---|---|
| | | CONTROL | CORT | LA 10 | LA 20 | FLUV |
| 1 | **Test for normal distribution** | | | | | |
| 2 | **Shapiro-Wilk test** | | | | | |
| 3 | W | 0.9108 | 0.8265 | 0.9728 | 0.8133 | 0.9481 |
| 4 | P value | 0.4414 | 0.1003 | 0.9109 | 0.0772 | 0.7252 |
| 5 | Passed normality test (alpha=0.05)? | Yes | Yes | Yes | Yes | Yes |
| 6 | P value summary | ns | ns | ns | ns | ns |
| 7 | | | | | | |
| 8 | **Number of values** | 6 | 6 | 6 | 6 | 6 |

| Ordinary one-way ANOVA<br>ANOVA results | | | | | |
|---|---|---|---|---|---|
| 1 | Table Analyzed | TBARS CORPO ESTRIAD | | | |
| 2 | Data sets analyzed | A-E | | | |
| 3 | | | | | |
| 4 | **ANOVA summary** | | | | |
| 5 | F | 3.101 | | | |
| 6 | P value | 0.0334 | | | |
| 7 | P value summary | * | | | |
| 8 | Significant diff. among means (P < 0.05)? | Yes | | | |
| 9 | R squared | 0.3316 | | | |
| 10 | | | | | |
| 11 | **Brown-Forsythe test** | | | | |
| 12 | F (DFn, DFd) | 3.457 (4, 25) | | | |
| 13 | P value | 0.0222 | | | |
| 14 | P value summary | * | | | |
| 15 | Are SDs significantly different (P < 0.05)? | Yes | | | |
| 16 | | | | | |
| 17 | **Bartlett's test** | | | | |
| 18 | Bartlett's statistic (corrected) | 34.39 | | | |
| 19 | P value | <0.0001 | | | |
| 20 | P value summary | **** | | | |
| 21 | Are SDs significantly different (P < 0.05)? | Yes | | | |
| 22 | | | | | |
| 23 | **ANOVA table** | **SS** | **DF** | **MS** | **F (DFn, DFd)** |
| 24 | Treatment (between columns) | 11293315 | 4 | 2823329 | F (4, 25) = 3.101 |
| 25 | Residual (within columns) | 22760023 | 25 | 910401 | |
| 26 | Total | 34053338 | 29 | | |
| 27 | | | | | |
| 28 | **Data summary** | | | | |
| 29 | Number of treatments (columns) | 5 | | | |
| 30 | Number of values (total) | 30 | | | |

| | |
|---|---|
| 1 | |
| 2 | |
| 3 | |
| 4 | |
| 5 | |
| 6 | |
| 7 | |
| 8 | |
| 9 | |
| 10 | |
| 11 | |
| 12 | |
| 13 | |
| 14 | |
| 15 | |
| 16 | |
| 17 | |
| 18 | |
| 19 | |
| 20 | |
| 21 | |
| 22 | |
| 23 | **P value** |
| 24 | P=0.0334 |
| 25 | |
| 26 | |
| 27 | |
| 28 | |
| 29 | |
| 30 | |

| **Ordinary one-way ANOVA** Multiple comparisons | | | | | |
|---|---|---|---|---|---|
| 1 | Number of families | 1 | | | |
| 2 | Number of comparisons per family | 10 | | | |
| 3 | Alpha | 0.05 | | | |
| 4 | | | | | |
| 5 | **Tukey's multiple comparisons test** | **Mean Diff.** | **95.00% CI of diff.** | **Significant?** | **Summary** |
| 6 | CONTROL vs. CORT | -1560 | -3178 to 58.03 | No | ns |
| 7 | CONTROL vs. LA 10 | -31.00 | -1649 to 1587 | No | ns |
| 8 | CONTROL vs. LA 20 | 34.50 | -1583 to 1652 | No | ns |
| 9 | CONTROL vs. FLUV | -645.3 | -2263 to 972.5 | No | ns |
| 10 | CORT vs. LA 10 | 1529 | -89.03 to 3147 | No | ns |
| 11 | CORT vs. LA 20 | 1594 | -23.53 to 3212 | No | ns |
| 12 | CORT vs. FLUV | 914.5 | -703.4 to 2532 | No | ns |
| 13 | LA 10 vs. LA 20 | 65.50 | -1552 to 1683 | No | ns |
| 14 | LA 10 vs. FLUV | -614.3 | -2232 to 1004 | No | ns |
| 15 | LA 20 vs. FLUV | -679.8 | -2298 to 938.0 | No | ns |
| 16 | | | | | |
| 17 | **Test details** | **Mean 1** | **Mean 2** | **Mean Diff.** | **SE of diff.** |
| 18 | CONTROL vs. CORT | 670.8 | 2231 | -1560 | 550.9 |
| 19 | CONTROL vs. LA 10 | 670.8 | 701.8 | -31.00 | 550.9 |
| 20 | CONTROL vs. LA 20 | 670.8 | 636.3 | 34.50 | 550.9 |
| 21 | CONTROL vs. FLUV | 670.8 | 1316 | -645.3 | 550.9 |
| 22 | CORT vs. LA 10 | 2231 | 701.8 | 1529 | 550.9 |
| 23 | CORT vs. LA 20 | 2231 | 636.3 | 1594 | 550.9 |
| 24 | CORT vs. FLUV | 2231 | 1316 | 914.5 | 550.9 |
| 25 | LA 10 vs. LA 20 | 701.8 | 636.3 | 65.50 | 550.9 |
| 26 | LA 10 vs. FLUV | 701.8 | 1316 | -614.3 | 550.9 |
| 27 | LA 20 vs. FLUV | 636.3 | 1316 | -679.8 | 550.9 |

| | | | | |
|---|---|---|---|---|
| 1 | | | | |
| 2 | | | | |
| 3 | | | | |
| 4 | | | | |
| 5 | **Adjusted P Value** | | | |
| 6 | 0.0627 | A-B | | |
| 7 | >0.9999 | A-C | | |
| 8 | >0.9999 | A-D | | |
| 9 | 0.7671 | A-E | | |
| 10 | 0.0705 | B-C | | |
| 11 | 0.0548 | B-D | | |
| 12 | 0.4753 | B-E | | |
| 13 | >0.9999 | C-D | | |
| 14 | 0.7971 | C-E | | |
| 15 | 0.7320 | D-E | | |
| 16 | | | | |
| 17 | **n1** | **n2** | **q** | **DF** |
| 18 | 6 | 6 | 4.004 | 25 |
| 19 | 6 | 6 | 0.07958 | 25 |
| 20 | 6 | 6 | 0.08857 | 25 |
| 21 | 6 | 6 | 1.657 | 25 |
| 22 | 6 | 6 | 3.925 | 25 |
| 23 | 6 | 6 | 4.093 | 25 |
| 24 | 6 | 6 | 2.348 | 25 |
| 25 | 6 | 6 | 0.1682 | 25 |
| 26 | 6 | 6 | 1.577 | 25 |
| 27 | 6 | 6 | 1.745 | 25 |

| Normality and Lognormality Tests | | A | B | C | D | E |
|---|---|---|---|---|---|---|
| | | CONTROL | CORT | LA 10 | LA 20 | FLUV |
| 1 | **Test for normal distribution** | | | | | |
| 2 | **Shapiro-Wilk test** | | | | | |
| 3 | W | 0.9199 | 0.6029 | 0.9382 | 0.9291 | 0.8830 |
| 4 | P value | 0.5043 | 0.0005 | 0.6448 | 0.5729 | 0.2831 |
| 5 | Passed normality test (alpha=0.05)? | Yes | No | Yes | Yes | Yes |
| 6 | P value summary | ns | *** | ns | ns | ns |
| 7 | | | | | | |
| 8 | **Number of values** | 6 | 6 | 6 | 6 | 6 |

| **Kruskal-Wallis test**<br>ANOVA results | | |
|---|---|---|
| 1 | Table Analyzed | Nitrito Hipocampo |
| 2 | | |
| 3 | **Kruskal-Wallis test** | |
| 4 | P value | 0.0758 |
| 5 | Exact or approximate P value? | Approximate |
| 6 | P value summary | ns |
| 7 | Do the medians vary signif. (P < 0.05)? | No |
| 8 | Number of groups | 5 |
| 9 | Kruskal-Wallis statistic | 8.469 |
| 10 | | |
| 11 | **Data summary** | |
| 12 | Number of treatments (columns) | 5 |
| 13 | Number of values (total) | 30 |

| | **Kruskal-Wallis test**<br>Multiple comparisons | | | | |
|---|---|---|---|---|---|
| 1 | Number of families | 1 | | | |
| 2 | Number of comparisons per family | 10 | | | |
| 3 | Alpha | 0.05 | | | |
| 4 | | | | | |
| 5 | **Dunn's multiple comparisons test** | **Mean rank diff.** | **Significant?** | **Summary** | **Adjusted P Value** |
| 6 | CONTROL vs. CORT | -2.250 | No | ns | >0.9999 |
| 7 | CONTROL vs. LA 10 | 7.333 | No | ns | >0.9999 |
| 8 | CONTROL vs. LA 20 | 8.917 | No | ns | 0.7931 |
| 9 | CONTROL vs. FLUV | -1.500 | No | ns | >0.9999 |
| 10 | CORT vs. LA 10 | 9.583 | No | ns | 0.5931 |
| 11 | CORT vs. LA 20 | 11.17 | No | ns | 0.2798 |
| 12 | CORT vs. FLUV | 0.7500 | No | ns | >0.9999 |
| 13 | LA 10 vs. LA 20 | 1.583 | No | ns | >0.9999 |
| 14 | LA 10 vs. FLUV | -8.833 | No | ns | 0.8215 |
| 15 | LA 20 vs. FLUV | -10.42 | No | ns | 0.4037 |
| 16 | | | | | |
| 17 | **Test details** | **Mean rank 1** | **Mean rank 2** | **Mean rank diff.** | **n1** |
| 18 | CONTROL vs. CORT | 18.00 | 20.25 | -2.250 | 6 |
| 19 | CONTROL vs. LA 10 | 18.00 | 10.67 | 7.333 | 6 |
| 20 | CONTROL vs. LA 20 | 18.00 | 9.083 | 8.917 | 6 |
| 21 | CONTROL vs. FLUV | 18.00 | 19.50 | -1.500 | 6 |
| 22 | CORT vs. LA 10 | 20.25 | 10.67 | 9.583 | 6 |
| 23 | CORT vs. LA 20 | 20.25 | 9.083 | 11.17 | 6 |
| 24 | CORT vs. FLUV | 20.25 | 19.50 | 0.7500 | 6 |
| 25 | LA 10 vs. LA 20 | 10.67 | 9.083 | 1.583 | 6 |
| 26 | LA 10 vs. FLUV | 10.67 | 19.50 | -8.833 | 6 |
| 27 | LA 20 vs. FLUV | 9.083 | 19.50 | -10.42 | 6 |

| | | |
|---|---|---|
| 1 | | |
| 2 | | |
| 3 | | |
| 4 | | |
| 5 | | |
| 6 | A-B | |
| 7 | A-C | |
| 8 | A-D | |
| 9 | A-E | |
| 10 | B-C | |
| 11 | B-D | |
| 12 | B-E | |
| 13 | C-D | |
| 14 | C-E | |
| 15 | D-E | |
| 16 | | |
| 17 | **n2** | **Z** |
| 18 | 6 | 0.4428 |
| 19 | 6 | 1.443 |
| 20 | 6 | 1.755 |
| 21 | 6 | 0.2952 |
| 22 | 6 | 1.886 |
| 23 | 6 | 2.198 |
| 24 | 6 | 0.1476 |
| 25 | 6 | 0.3116 |
| 26 | 6 | 1.738 |
| 27 | 6 | 2.050 |

| Normality and Lognormality Tests | | A | B | C | D | E |
|---|---|---|---|---|---|---|
| | | CONTROL | CORT | LA 10 | LA 20 | FLUV |
| 1 | **Test for normal distribution** | | | | | |
| 2 | **Shapiro-Wilk test** | | | | | |
| 3 | W | 0.9777 | 0.9225 | 0.9584 | 0.9643 | 0.9126 |
| 4 | P value | 0.9394 | 0.5239 | 0.8074 | 0.8525 | 0.4537 |
| 5 | Passed normality test (alpha=0.05)? | Yes | Yes | Yes | Yes | Yes |
| 6 | P value summary | ns | ns | ns | ns | ns |
| 7 | | | | | | |
| 8 | **Number of values** | 6 | 6 | 6 | 6 | 6 |

| Ordinary one-way ANOVA<br>ANOVA results | | | | | |
|---|---|---|---|---|---|
| 1 | Table Analyzed | Nitrito CÓRTEX PRÉ-FRONTAL | | | |
| 2 | Data sets analyzed | A-E | | | |
| 3 | | | | | |
| 4 | **ANOVA summary** | | | | |
| 5 | F | 0.4778 | | | |
| 6 | P value | 0.7517 | | | |
| 7 | P value summary | ns | | | |
| 8 | Significant diff. among means (P < 0.05)? | No | | | |
| 9 | R squared | 0.07102 | | | |
| 10 | | | | | |
| 11 | **Brown-Forsythe test** | | | | |
| 12 | F (DFn, DFd) | 0.5466 (4, 25) | | | |
| 13 | P value | 0.7031 | | | |
| 14 | P value summary | ns | | | |
| 15 | Are SDs significantly different (P < 0.05)? | No | | | |
| 16 | | | | | |
| 17 | **Bartlett's test** | | | | |
| 18 | Bartlett's statistic (corrected) | 3.627 | | | |
| 19 | P value | 0.4589 | | | |
| 20 | P value summary | ns | | | |
| 21 | Are SDs significantly different (P < 0.05)? | No | | | |
| 22 | | | | | |
| 23 | **ANOVA table** | **SS** | **DF** | **MS** | **F (DFn, DFd)** |
| 24 | Treatment (between columns) | 6.722 | 4 | 1.681 | F (4, 25) = 0.4778 |
| 25 | Residual (within columns) | 87.93 | 25 | 3.517 | |
| 26 | Total | 94.65 | 29 | | |
| 27 | | | | | |
| 28 | **Data summary** | | | | |
| 29 | Number of treatments (columns) | 5 | | | |
| 30 | Number of values (total) | 30 | | | |

| | |
|---|---|
| 1 | |
| 2 | |
| 3 | |
| 4 | |
| 5 | |
| 6 | |
| 7 | |
| 8 | |
| 9 | |
| 10 | |
| 11 | |
| 12 | |
| 13 | |
| 14 | |
| 15 | |
| 16 | |
| 17 | |
| 18 | |
| 19 | |
| 20 | |
| 21 | |
| 22 | |
| 23 | **P value** |
| 24 | P=0.7517 |
| 25 | |
| 26 | |
| 27 | |
| 28 | |
| 29 | |
| 30 | |

| Ordinary one-way ANOVA<br>Multiple comparisons | | | | | |
|---|---|---|---|---|---|
| 1 | Number of families | 1 | | | |
| 2 | Number of comparisons per family | 10 | | | |
| 3 | Alpha | 0.05 | | | |
| 4 | | | | | |
| 5 | **Tukey's multiple comparisons test** | **Mean Diff.** | **95.00% CI of diff.** | **Significant?** | **Summary** |
| 6 | CONTROL vs. CORT | -0.5693 | -3.749 to 2.611 | No | ns |
| 7 | CONTROL vs. LA 10 | 0.7064 | -2.474 to 3.886 | No | ns |
| 8 | CONTROL vs. LA 20 | -0.5281 | -3.708 to 2.652 | No | ns |
| 9 | CONTROL vs. FLUV | -0.3635 | -3.544 to 2.816 | No | ns |
| 10 | CORT vs. LA 10 | 1.276 | -1.904 to 4.456 | No | ns |
| 11 | CORT vs. LA 20 | 0.04115 | -3.139 to 3.221 | No | ns |
| 12 | CORT vs. FLUV | 0.2058 | -2.974 to 3.386 | No | ns |
| 13 | LA 10 vs. LA 20 | -1.235 | -4.415 to 1.945 | No | ns |
| 14 | LA 10 vs. FLUV | -1.070 | -4.250 to 2.110 | No | ns |
| 15 | LA 20 vs. FLUV | 0.1646 | -3.015 to 3.345 | No | ns |
| 16 | | | | | |
| 17 | **Test details** | **Mean 1** | **Mean 2** | **Mean Diff.** | **SE of diff.** |
| 18 | CONTROL vs. CORT | 3.768 | 4.337 | -0.5693 | 1.083 |
| 19 | CONTROL vs. LA 10 | 3.768 | 3.062 | 0.7064 | 1.083 |
| 20 | CONTROL vs. LA 20 | 3.768 | 4.296 | -0.5281 | 1.083 |
| 21 | CONTROL vs. FLUV | 3.768 | 4.132 | -0.3635 | 1.083 |
| 22 | CORT vs. LA 10 | 4.337 | 3.062 | 1.276 | 1.083 |
| 23 | CORT vs. LA 20 | 4.337 | 4.296 | 0.04115 | 1.083 |
| 24 | CORT vs. FLUV | 4.337 | 4.132 | 0.2058 | 1.083 |
| 25 | LA 10 vs. LA 20 | 3.062 | 4.296 | -1.235 | 1.083 |
| 26 | LA 10 vs. FLUV | 3.062 | 4.132 | -1.070 | 1.083 |
| 27 | LA 20 vs. FLUV | 4.296 | 4.132 | 0.1646 | 1.083 |

| | | | | |
|---|---|---|---|---|
| 1 | | | | |
| 2 | | | | |
| 3 | | | | |
| 4 | | | | |
| 5 | **Adjusted P Value** | | | |
| 6 | 0.9839 | A-B | | |
| 7 | 0.9646 | A-C | | |
| 8 | 0.9878 | A-D | | |
| 9 | 0.9971 | A-E | | |
| 10 | 0.7634 | B-C | | |
| 11 | >0.9999 | B-D | | |
| 12 | 0.9997 | B-E | | |
| 13 | 0.7840 | C-D | | |
| 14 | 0.8581 | C-E | | |
| 15 | 0.9999 | D-E | | |
| 16 | | | | |
| 17 | **n1** | **n2** | **q** | **DF** |
| 18 | 6 | 6 | 0.7435 | 25 |
| 19 | 6 | 6 | 0.9227 | 25 |
| 20 | 6 | 6 | 0.6898 | 25 |
| 21 | 6 | 6 | 0.4748 | 25 |
| 22 | 6 | 6 | 1.666 | 25 |
| 23 | 6 | 6 | 0.05375 | 25 |
| 24 | 6 | 6 | 0.2687 | 25 |
| 25 | 6 | 6 | 1.612 | 25 |
| 26 | 6 | 6 | 1.397 | 25 |
| 27 | 6 | 6 | 0.2150 | 25 |

| Normality and Lognormality Tests | | A<br>CONTROL | B<br>CORT | C<br>LA 10 | D<br>LA 20 | E<br>FLUV |
|---|---|---|---|---|---|---|
| 1 | **Test for normal distribution** | | | | | |
| 2 | **Shapiro-Wilk test** | | | | | |
| 3 | W | 0.7579 | 0.8598 | 0.8391 | 0.8209 | 0.8351 |
| 4 | P value | 0.0238 | 0.1885 | 0.1281 | 0.0899 | 0.1186 |
| 5 | Passed normality test (alpha=0.05)? | No | Yes | Yes | Yes | Yes |
| 6 | P value summary | * | ns | ns | ns | ns |
| 7 | | | | | | |
| 8 | **Number of values** | 6 | 6 | 6 | 6 | 6 |

| **Kruskal-Wallis test**<br>ANOVA results | | |
|---|---|---|
| 1 | Table Analyzed | Nitrito CORPO ESTRIADO |
| 2 | | |
| 3 | **Kruskal-Wallis test** | |
| 4 | P value | 0.0045 |
| 5 | Exact or approximate P value? | Approximate |
| 6 | P value summary | ** |
| 7 | Do the medians vary signif. (P < 0.05)? | Yes |
| 8 | Number of groups | 5 |
| 9 | Kruskal-Wallis statistic | 15.12 |
| 10 | | |
| 11 | **Data summary** | |
| 12 | Number of treatments (columns) | 5 |
| 13 | Number of values (total) | 30 |

| **Kruskal-Wallis test** Multiple comparisons | | | | | |
|---|---|---|---|---|---|
| 1 | Number of families | 1 | | | |
| 2 | Number of comparisons per family | 10 | | | |
| 3 | Alpha | 0.05 | | | |
| 4 | | | | | |
| 5 | **Dunn's multiple comparisons test** | **Mean rank diff.** | **Significant?** | **Summary** | **Adjusted P Value** |
| 6 | CONTROL vs. CORT | -11.17 | No | ns | 0.2797 |
| 7 | CONTROL vs. LA 10 | 8.000 | No | ns | >0.9999 |
| 8 | CONTROL vs. LA 20 | -2.167 | No | ns | >0.9999 |
| 9 | CONTROL vs. FLUV | -4.667 | No | ns | >0.9999 |
| 10 | CORT vs. LA 10 | 19.17 | Yes | ** | 0.0016 |
| 11 | CORT vs. LA 20 | 9.000 | No | ns | 0.7651 |
| 12 | CORT vs. FLUV | 6.500 | No | ns | >0.9999 |
| 13 | LA 10 vs. LA 20 | -10.17 | No | ns | 0.4540 |
| 14 | LA 10 vs. FLUV | -12.67 | No | ns | 0.1267 |
| 15 | LA 20 vs. FLUV | -2.500 | No | ns | >0.9999 |
| 16 | | | | | |
| 17 | **Test details** | **Mean rank 1** | **Mean rank 2** | **Mean rank diff.** | **n1** |
| 18 | CONTROL vs. CORT | 13.50 | 24.67 | -11.17 | 6 |
| 19 | CONTROL vs. LA 10 | 13.50 | 5.500 | 8.000 | 6 |
| 20 | CONTROL vs. LA 20 | 13.50 | 15.67 | -2.167 | 6 |
| 21 | CONTROL vs. FLUV | 13.50 | 18.17 | -4.667 | 6 |
| 22 | CORT vs. LA 10 | 24.67 | 5.500 | 19.17 | 6 |
| 23 | CORT vs. LA 20 | 24.67 | 15.67 | 9.000 | 6 |
| 24 | CORT vs. FLUV | 24.67 | 18.17 | 6.500 | 6 |
| 25 | LA 10 vs. LA 20 | 5.500 | 15.67 | -10.17 | 6 |
| 26 | LA 10 vs. FLUV | 5.500 | 18.17 | -12.67 | 6 |
| 27 | LA 20 vs. FLUV | 15.67 | 18.17 | -2.500 | 6 |

| | | |
|---|---|---|
| 1 | | |
| 2 | | |
| 3 | | |
| 4 | | |
| 5 | | |
| 6 | A-B | |
| 7 | A-C | |
| 8 | A-D | |
| 9 | A-E | |
| 10 | B-C | |
| 11 | B-D | |
| 12 | B-E | |
| 13 | C-D | |
| 14 | C-E | |
| 15 | D-E | |
| 16 | | |
| 17 | **n2** | **Z** |
| 18 | 6 | 2.198 |
| 19 | 6 | 1.575 |
| 20 | 6 | 0.4264 |
| 21 | 6 | 0.9185 |
| 22 | 6 | 3.772 |
| 23 | 6 | 1.771 |
| 24 | 6 | 1.279 |
| 25 | 6 | 2.001 |
| 26 | 6 | 2.493 |
| 27 | 6 | 0.4920 |

| Normality and Lognormality Tests | | A | B | C | D | E |
|---|---|---|---|---|---|---|
| | | CONTROL | CORT | LA 10 | LA 20 | FLUV |
| 1 | **Test for normal distribution** | | | | | |
| 2 | **Shapiro-Wilk test** | | | | | |
| 3 | W | 0.7582 | 0.9275 | 0.9277 | 0.7561 | 0.9869 |
| 4 | P value | 0.0239 | 0.5613 | 0.5622 | 0.0229 | 0.9802 |
| 5 | Passed normality test (alpha=0.05)? | No | Yes | Yes | No | Yes |
| 6 | P value summary | * | ns | ns | * | ns |
| 7 | | | | | | |
| 8 | **Number of values** | 6 | 6 | 6 | 6 | 6 |

| Kruskal-Wallis test<br>ANOVA results | | |
|---|---|---|
| 1 | Table Analyzed | GSH - HIPOCAMP( |
| 2 | | |
| 3 | **Kruskal-Wallis test** | |
| 4 | P value | 0.2361 |
| 5 | Exact or approximate P value? | Approximate |
| 6 | P value summary | ns |
| 7 | Do the medians vary signif. (P < 0.05)? | No |
| 8 | Number of groups | 5 |
| 9 | Kruskal-Wallis statistic | 5.542 |
| 10 | | |
| 11 | **Data summary** | |
| 12 | Number of treatments (columns) | 5 |
| 13 | Number of values (total) | 30 |

| **Kruskal-Wallis test**<br>Multiple comparisons | | | | | |
|---|---|---|---|---|---|
| 1 | Number of families | 1 | | | |
| 2 | Number of comparisons per family | 10 | | | |
| 3 | Alpha | 0.05 | | | |
| 4 | | | | | |
| 5 | **Dunn's multiple comparisons test** | **Mean rank diff.** | **Significant?** | **Summary** | **Adjusted P Value** |
| 6 | CONTROL vs. CORT | 3.833 | No | ns | >0.9999 |
| 7 | CONTROL vs. LA 10 | -4.333 | No | ns | >0.9999 |
| 8 | CONTROL vs. LA 20 | 6.250 | No | ns | >0.9999 |
| 9 | CONTROL vs. FLUV | -1.583 | No | ns | >0.9999 |
| 10 | CORT vs. LA 10 | -8.167 | No | ns | >0.9999 |
| 11 | CORT vs. LA 20 | 2.417 | No | ns | >0.9999 |
| 12 | CORT vs. FLUV | -5.417 | No | ns | >0.9999 |
| 13 | LA 10 vs. LA 20 | 10.58 | No | ns | 0.3730 |
| 14 | LA 10 vs. FLUV | 2.750 | No | ns | >0.9999 |
| 15 | LA 20 vs. FLUV | -7.833 | No | ns | >0.9999 |
| 16 | | | | | |
| 17 | **Test details** | **Mean rank 1** | **Mean rank 2** | **Mean rank diff.** | **n1** |
| 18 | CONTROL vs. CORT | 16.33 | 12.50 | 3.833 | 6 |
| 19 | CONTROL vs. LA 10 | 16.33 | 20.67 | -4.333 | 6 |
| 20 | CONTROL vs. LA 20 | 16.33 | 10.08 | 6.250 | 6 |
| 21 | CONTROL vs. FLUV | 16.33 | 17.92 | -1.583 | 6 |
| 22 | CORT vs. LA 10 | 12.50 | 20.67 | -8.167 | 6 |
| 23 | CORT vs. LA 20 | 12.50 | 10.08 | 2.417 | 6 |
| 24 | CORT vs. FLUV | 12.50 | 17.92 | -5.417 | 6 |
| 25 | LA 10 vs. LA 20 | 20.67 | 10.08 | 10.58 | 6 |
| 26 | LA 10 vs. FLUV | 20.67 | 17.92 | 2.750 | 6 |
| 27 | LA 20 vs. FLUV | 10.08 | 17.92 | -7.833 | 6 |

| | | |
|---|---|---|
| 1 | | |
| 2 | | |
| 3 | | |
| 4 | | |
| 5 | | |
| 6 | A-B | |
| 7 | A-C | |
| 8 | A-D | |
| 9 | A-E | |
| 10 | B-C | |
| 11 | B-D | |
| 12 | B-E | |
| 13 | C-D | |
| 14 | C-E | |
| 15 | D-E | |
| 16 | | |
| 17 | **n2** | **Z** |
| 18 | 6 | 0.7543 |
| 19 | 6 | 0.8527 |
| 20 | 6 | 1.230 |
| 21 | 6 | 0.3116 |
| 22 | 6 | 1.607 |
| 23 | 6 | 0.4755 |
| 24 | 6 | 1.066 |
| 25 | 6 | 2.082 |
| 26 | 6 | 0.5411 |
| 27 | 6 | 1.541 |

| **Normality and Lognormality Tests**<br>Tabular results | | A<br>CONTROL | B<br>CORT | C<br>LA 10 | D<br>LA 20 | E<br>FLUV |
|---|---|---|---|---|---|---|
| 1 | **Test for normal distribution** | | | | | |
| 2 | **Shapiro-Wilk test** | | | | | |
| 3 | W | 0.8545 | 0.9158 | 0.8205 | 0.9484 | 0.9652 |
| 4 | P value | 0.1712 | 0.4755 | 0.0891 | 0.7274 | 0.8586 |
| 5 | Passed normality test (alpha=0.05)? | Yes | Yes | Yes | Yes | Yes |
| 6 | P value summary | ns | ns | ns | ns | ns |
| 7 | | | | | | |
| 8 | **Number of values** | 6 | 6 | 6 | 6 | 6 |

| Kruskal-Wallis test<br>ANOVA results | | |
|---|---|---|
| 1 | Table Analyzed | GSH - CÓRTEX PRÉ-FRONTA |
| 2 | | |
| 3 | **Kruskal-Wallis test** | |
| 4 | P value | 0.1038 |
| 5 | Exact or approximate P value? | Approximate |
| 6 | P value summary | ns |
| 7 | Do the medians vary signif. (P < 0.05)? | No |
| 8 | Number of groups | 5 |
| 9 | Kruskal-Wallis statistic | 7.686 |
| 10 | | |
| 11 | **Data summary** | |
| 12 | Number of treatments (columns) | 5 |
| 13 | Number of values (total) | 30 |

| **Kruskal-Wallis test**<br>Multiple comparisons | | | | | |
|---|---|---|---|---|---|
| 1 | Number of families | 1 | | | |
| 2 | Number of comparisons per family | 10 | | | |
| 3 | Alpha | 0.05 | | | |
| 4 | | | | | |
| 5 | **Dunn's multiple comparisons test** | **Mean rank diff.** | **Significant?** | **Summary** | **Adjusted P Value** |
| 6 | CONTROL vs. CORT | 1.500 | No | ns | >0.9999 |
| 7 | CONTROL vs. LA 10 | 11.83 | No | ns | 0.1990 |
| 8 | CONTROL vs. LA 20 | 8.167 | No | ns | >0.9999 |
| 9 | CONTROL vs. FLUV | 2.667 | No | ns | >0.9999 |
| 10 | CORT vs. LA 10 | 10.33 | No | ns | 0.4205 |
| 11 | CORT vs. LA 20 | 6.667 | No | ns | >0.9999 |
| 12 | CORT vs. FLUV | 1.167 | No | ns | >0.9999 |
| 13 | LA 10 vs. LA 20 | -3.667 | No | ns | >0.9999 |
| 14 | LA 10 vs. FLUV | -9.167 | No | ns | 0.7131 |
| 15 | LA 20 vs. FLUV | -5.500 | No | ns | >0.9999 |
| 16 | | | | | |
| 17 | **Test details** | **Mean rank 1** | **Mean rank 2** | **Mean rank diff.** | **n1** |
| 18 | CONTROL vs. CORT | 20.33 | 18.83 | 1.500 | 6 |
| 19 | CONTROL vs. LA 10 | 20.33 | 8.500 | 11.83 | 6 |
| 20 | CONTROL vs. LA 20 | 20.33 | 12.17 | 8.167 | 6 |
| 21 | CONTROL vs. FLUV | 20.33 | 17.67 | 2.667 | 6 |
| 22 | CORT vs. LA 10 | 18.83 | 8.500 | 10.33 | 6 |
| 23 | CORT vs. LA 20 | 18.83 | 12.17 | 6.667 | 6 |
| 24 | CORT vs. FLUV | 18.83 | 17.67 | 1.167 | 6 |
| 25 | LA 10 vs. LA 20 | 8.500 | 12.17 | -3.667 | 6 |
| 26 | LA 10 vs. FLUV | 8.500 | 17.67 | -9.167 | 6 |
| 27 | LA 20 vs. FLUV | 12.17 | 17.67 | -5.500 | 6 |

| | | |
|---|---|---|
| 1 | | |
| 2 | | |
| 3 | | |
| 4 | | |
| 5 | | |
| 6 | A-B | |
| 7 | A-C | |
| 8 | A-D | |
| 9 | A-E | |
| 10 | B-C | |
| 11 | B-D | |
| 12 | B-E | |
| 13 | C-D | |
| 14 | C-E | |
| 15 | D-E | |
| 16 | | |
| 17 | **n2** | **Z** |
| 18 | 6 | 0.2951 |
| 19 | 6 | 2.328 |
| 20 | 6 | 1.607 |
| 21 | 6 | 0.5247 |
| 22 | 6 | 2.033 |
| 23 | 6 | 1.312 |
| 24 | 6 | 0.2295 |
| 25 | 6 | 0.7214 |
| 26 | 6 | 1.804 |
| 27 | 6 | 1.082 |

| Normality and Lognormality Tests | | A | B | C | D | E |
|---|---|---|---|---|---|---|
| | | CONTROL | CORT | LA 10 | LA 20 | FLUV |
| 1 | **Test for normal distribution** | | | | | |
| 2 | **Shapiro-Wilk test** | | | | | |
| 3 | W | 0.7765 | 0.9348 | 0.8623 | 0.9025 | 0.9533 |
| 4 | P value | 0.0358 | 0.6174 | 0.1973 | 0.3887 | 0.7673 |
| 5 | Passed normality test (alpha=0.05)? | No | Yes | Yes | Yes | Yes |
| 6 | P value summary | * | ns | ns | ns | ns |
| 7 | | | | | | |
| 8 | **Number of values** | 6 | 6 | 6 | 6 | 6 |

| Kruskal-Wallis test<br>ANOVA results | | |
|---|---|---|
| 1 | Table Analyzed | GSH - CORPO ESTRIADO |
| 2 | | |
| 3 | **Kruskal-Wallis test** | |
| 4 | P value | 0.1272 |
| 5 | Exact or approximate P value? | Approximate |
| 6 | P value summary | ns |
| 7 | Do the medians vary signif. (P < 0.05)? | No |
| 8 | Number of groups | 5 |
| 9 | Kruskal-Wallis statistic | 7.170 |
| 10 | | |
| 11 | **Data summary** | |
| 12 | Number of treatments (columns) | 5 |
| 13 | Number of values (total) | 30 |

| Kruskal-Wallis test<br>Multiple comparisons | | | | | |
|---|---|---|---|---|---|
| 1 | Number of families | 1 | | | |
| 2 | Number of comparisons per family | 10 | | | |
| 3 | Alpha | 0.05 | | | |
| 4 | | | | | |
| 5 | **Dunn's multiple comparisons test** | **Mean rank diff.** | **Significant?** | **Summary** | **Adjusted P Value** |
| 6 | CONTROL vs. CORT | -0.7500 | No | ns | >0.9999 |
| 7 | CONTROL vs. LA 10 | 5.333 | No | ns | >0.9999 |
| 8 | CONTROL vs. LA 20 | 10.17 | No | ns | 0.4542 |
| 9 | CONTROL vs. FLUV | -0.5833 | No | ns | >0.9999 |
| 10 | CORT vs. LA 10 | 6.083 | No | ns | >0.9999 |
| 11 | CORT vs. LA 20 | 10.92 | No | ns | 0.3169 |
| 12 | CORT vs. FLUV | 0.1667 | No | ns | >0.9999 |
| 13 | LA 10 vs. LA 20 | 4.833 | No | ns | >0.9999 |
| 14 | LA 10 vs. FLUV | -5.917 | No | ns | >0.9999 |
| 15 | LA 20 vs. FLUV | -10.75 | No | ns | 0.3439 |
| 16 | | | | | |
| 17 | **Test details** | **Mean rank 1** | **Mean rank 2** | **Mean rank diff.** | **n1** |
| 18 | CONTROL vs. CORT | 18.33 | 19.08 | -0.7500 | 6 |
| 19 | CONTROL vs. LA 10 | 18.33 | 13.00 | 5.333 | 6 |
| 20 | CONTROL vs. LA 20 | 18.33 | 8.167 | 10.17 | 6 |
| 21 | CONTROL vs. FLUV | 18.33 | 18.92 | -0.5833 | 6 |
| 22 | CORT vs. LA 10 | 19.08 | 13.00 | 6.083 | 6 |
| 23 | CORT vs. LA 20 | 19.08 | 8.167 | 10.92 | 6 |
| 24 | CORT vs. FLUV | 19.08 | 18.92 | 0.1667 | 6 |
| 25 | LA 10 vs. LA 20 | 13.00 | 8.167 | 4.833 | 6 |
| 26 | LA 10 vs. FLUV | 13.00 | 18.92 | -5.917 | 6 |
| 27 | LA 20 vs. FLUV | 8.167 | 18.92 | -10.75 | 6 |

| | | |
|---|---|---|
| 1 | | |
| 2 | | |
| 3 | | |
| 4 | | |
| 5 | | |
| 6 | A-B | |
| 7 | A-C | |
| 8 | A-D | |
| 9 | A-E | |
| 10 | B-C | |
| 11 | B-D | |
| 12 | B-E | |
| 13 | C-D | |
| 14 | C-E | |
| 15 | D-E | |
| 16 | | |
| 17 | **n2** | **Z** |
| 18 | 6 | 0.1476 |
| 19 | 6 | 1.050 |
| 20 | 6 | 2.001 |
| 21 | 6 | 0.1148 |
| 22 | 6 | 1.197 |
| 23 | 6 | 2.148 |
| 24 | 6 | 0.03280 |
| 25 | 6 | 0.9512 |
| 26 | 6 | 1.164 |
| 27 | 6 | 2.116 |

| Normality and Lognormality Tests | | A | B | C | D |
|---|---|---|---|---|---|
| | | Control | DIA 1 | DIA 14 | DIA 20 |
| 1 | **Test for normal distribution** | | | | |
| 2 | **Shapiro-Wilk test** | | | | |
| 3 | W | 0.7502 | 0.7627 | 0.9465 | 0.9130 |
| 4 | P value | 0.0003 | 0.0005 | 0.3731 | 0.0973 |
| 5 | Passed normality test (alpha=0.05)? | No | No | Yes | Yes |
| 6 | P value summary | *** | *** | ns | ns |
| 7 | | | | | |
| 8 | **Number of values** | 18 | 18 | 18 | 18 |

| Kruskal-Wallis test<br>ANOVA results | | |
|---|---|---|
| 1 | Table Analyzed | Evolução depressão |
| 2 | | |
| 3 | **Kruskal-Wallis test** | |
| 4 | P value | <0.0001 |
| 5 | Exact or approximate P value? | Approximate |
| 6 | P value summary | **** |
| 7 | Do the medians vary signif. (P < 0.05)? | Yes |
| 8 | Number of groups | 4 |
| 9 | Kruskal-Wallis statistic | 23.41 |
| 10 | | |
| 11 | **Data summary** | |
| 12 | Number of treatments (columns) | 4 |
| 13 | Number of values (total) | 72 |

| **Kruskal-Wallis test**<br>Multiple comparisons | | | | | |
|---|---|---|---|---|---|
| 1 | Number of families | 1 | | | |
| 2 | Number of comparisons per family | 6 | | | |
| 3 | Alpha | 0.05 | | | |
| 4 | | | | | |
| 5 | **Dunn's multiple comparisons test** | **Mean rank diff.** | **Significant?** | **Summary** | **Adjusted P Value** |
| 6 | Control vs. DIA 1 | -3.944 | No | ns | >0.9999 |
| 7 | Control vs. DIA 14 | -25.58 | Yes | ** | 0.0014 |
| 8 | Control vs. DIA 20 | -25.58 | Yes | ** | 0.0014 |
| 9 | DIA 1 vs. DIA 14 | -21.64 | Yes | * | 0.0111 |
| 10 | DIA 1 vs. DIA 20 | -21.64 | Yes | * | 0.0111 |
| 11 | DIA 14 vs. DIA 20 | 0.000 | No | ns | >0.9999 |
| 12 | | | | | |
| 13 | **Test details** | **Mean rank 1** | **Mean rank 2** | **Mean rank diff.** | **n1** |
| 14 | Control vs. DIA 1 | 22.72 | 26.67 | -3.944 | 18 |
| 15 | Control vs. DIA 14 | 22.72 | 48.31 | -25.58 | 18 |
| 16 | Control vs. DIA 20 | 22.72 | 48.31 | -25.58 | 18 |
| 17 | DIA 1 vs. DIA 14 | 26.67 | 48.31 | -21.64 | 18 |
| 18 | DIA 1 vs. DIA 20 | 26.67 | 48.31 | -21.64 | 18 |
| 19 | DIA 14 vs. DIA 20 | 48.31 | 48.31 | 0.000 | 18 |

| | | |
|---|---|---|
| 1 | | |
| 2 | | |
| 3 | | |
| 4 | | |
| 5 | | |
| 6 | A-B | |
| 7 | A-C | |
| 8 | A-D | |
| 9 | B-C | |
| 10 | B-D | |
| 11 | C-D | |
| 12 | | |
| 13 | **n2** | **Z** |
| 14 | 18 | 0.5676 |
| 15 | 18 | 3.681 |
| 16 | 18 | 3.681 |
| 17 | 18 | 3.114 |
| 18 | 18 | 3.114 |
| 19 | 18 | 0.000 |
